# Broad-spectrum extracellular antiviral properties of Cucurbit[n]urils

**DOI:** 10.1101/2022.03.15.484424

**Authors:** Luke M. Jones, Elana H. Super, Lauren J. Batt, Matteo Gasbarri, Benjamin T. Cheesman, Andrew M. Howe, Roger Coulston, Samuel T. Jones

**Affiliations:** Department of Materials and The Henry Royce Institute, The University of Manchester, Manchester, M19 3PL, UK; Institute of Materials, Interfaculty Bioengineering Institute, MXG 030, Lausanne, Switzerland; Aqdot Limited, Iconix Park, London Road, Pampisford, Cambridge, CB22 3EG, UK

## Abstract

Viruses are microscopic pathogens capable of causing disease and are responsible for a range of human mortality and morbidity worldwide. They can be rendered harmless or destroyed with a range of antiviral chemical compounds. Cucurbit[n]urils (CB[n]s) are a macrocycle chemical compound existing as a range of homologues; due to their structure they can bind to biological materials, acting as supramolecular “hosts” to “guests”, such as amino acids. Due to the increasing need for a non-toxic antiviral compound, we investigated whether cucurbit[n]urils could act in an antiviral manner. We have found that certain cucurbit[n]uril homologues do indeed have an antiviral effect against a range of viruses, including RSV and SARS-CoV-2. In particular, we demonstrate that CB[7] is the active homologue of CB[n] mixtures, having an antiviral effect against enveloped and non-enveloped species. High levels of efficacy were observed with five-minute contact times across different viruses. We also demonstrate that CB[7] acts with an extracellular virucidal mode of action via host-guest supramolecular interactions between viral surface proteins and the CB[n] cavity, rather than via cell internalisation or a virustatic mechanism. This finding demonstrates that CB[7] acts as a supramolecular virucidal antiviral (a mechanism distinct from other current extracellular antivirals) demonstrating the potential of supramolecular interactions for future antiviral disinfectants.

## MAIN TEXT

### Introduction

Viruses are harmful, microscopic pathogens that replicate within host cells before being spread via aerosols (coughs, sneezes), surface contamination, and/or infected biological fluids. The impact each has varies, but they can be responsible for severe disease in human hosts, and disease outbreaks (epidemics or pandemics), including the COVID-19 pandemic. Epidemics of viral diseases are also increasing in frequency [1, 2], with associated costs in overall mortality, morbidity and economic damage [3, 4]. Viruses themselves are composed of amino acids self-assembled into proteinaceous shells – capsids – protecting the genome. The capsid may or may not be surrounded by a lipid envelope. Regardless of envelope presence, all viruses sport attachment, or spike, proteins on their surface for the purpose of adhering to and infecting host cells [5].

To address the global threat posed by all viruses, there are three main interventions; vaccines, drugs or disinfection. Chemical disinfectants (such as bleach and soaps) are often broad-spectrum and have been deployed in a range of sprays, surface sanitisers, and hand-gels to prevent viral spread [6]. However, many of these substances are highly cytotoxic and reactive, meaning their safe applications can be limited. Less cytotoxic antiviral options have been previously studied, including quaternary ammonium compounds [7, 8], nanoparticles [9–11], and polymers [12–14]. Each works differently; some target the genetic material, some the viral capsid proteins, but all aim to inhibit the virus such that it is no longer infectious. However, many remain toxic and so their range of use is severely limited.

Macrocyclic compounds are frequently used as drug carriers [15, 16], including as carriers for antivirals [17, 18]. Additionally, due to their wide range of binding motifs, macrocycles offer the potential to interact directly with viruses and inhibit infection. However, to date the use of macrocycles as antivirals has been limited to modified macrocycles [19], or their use via indirect effects (for example via binding cholesterol [20]). Cucurbit[n]urils (CB[n]) are barrel-shaped macrocycles composed of ‘n’ glycoluril monomers/units linked by methylene bridges [21] that are produced in a mixture of ring sizes, which can then be separated into discrete sizes from n = 5-8 and 10 [22–24]. Like other macrocycles, CB[n]s have a cavity capable of supramolecular binding to a range of guest molecules [25], with the larger CBs (CB[8] and CB[10]) capable of inclusion of two guest simultaneously [26, 27]. This binding arises via guest interactions with the hydrophobic, unpolarisable cavity and polar, carbonyl-rich portals supported by the entropy gain on releasing water molecules from the cavity [22]. CBs have been widely studied and utilized to produce unique supramolecular structures for a range of applications in drug delivery [28, 29], tuning protein functionality [30] and forming hydrogels [31, 32]. CBs have also been shown to bind to an array of biologically relevant groups, including the amino acid constituents [33, 34] of proteins such as insulin [35]. Recently, it has been suggested that cucurbit[7]uril (CB[7]) is able to act as indirect antiviral by binding intracellularly to polyamines and disrupting the viral replication cycle [36].

We hypothosised that CB[n]s could bind directly to viruses via supramolecular interactions between exposed surface proteins and the CB cavity, which would then inhibit infection. This inhibition could be via reversible binding (virustatic) or may be irreversible if binding results in an irreversible conformational change (virucidal). Both possibilities would inhibit infection, but only a virucidal mechanism would lead to inactivation of the virus. This interaction is different to many other extracellular antivirals, which largely function by mimicking cell-binding regions [19, 37, 38] or leaching toxic metal ions [39, 40]. In order to test our hypotheses we used a range of viruses, culturable in vitro, and performed a number of different assays to determine the effectiveness and mode of action of the individual CB homologues CB[6], CB[7] and CB[8] in addition to the n = 6,7,8 mixture CB[n].

## Results

### Investigating Antiviral Efficacy

We began with solution-based testing to determine if any of our CB homologues exhibited antiviral efficacy. To investigate the efficacy of CB[n]s as antivirals we needed to identify an easily culturable model virus/cell system. For this we selected herpes simplex virus-2 (HSV-2) in Vero cells, which is a well-studied system across a range of assays, is easily cultured in-vitro and readily forms plaques in cell monolayers. In order to determine if CB[n]s were able to inhibit viral infection, median tissue culture infectious dose (TCID_50_) assays were used with HSV-2. The CB homologue mixture CB[n], as well as isolated CB[6], CB[7] and CB[8] were studied. Each CB sample was mixed with virus for 5 minutes before being serially diluted to determine viral titre. Initial TCID_50_ assays indicated that both CB[n] and CB[7] were effective at inhibiting HSV-2, with an approximately 5-log reduction in viral titre observed at 20-50 mg/ml. The lack of viral titre reduction observed in conjunction with CB[6] and CB[8] indicated that it was the CB[7] component of the CB[n] responsible for the reduction (Figure 1B). CB[n] and CB[7] were both significantly different from CB[6] and CB[8] (all adjusted p values <0.0001). CB[7] is more soluble in aqueous systems than CB[6] and CB[8] [41, 42], meaning more will be available to bind to the virus. In addition, the cavity of CB[6] is too small for binding to amino acids [43]. Conversely, the cavity of CB[8] is large enough to bind to, and has an affinity for, particular amino acids [44]. More experiments would be needed to determine why CB[8] lacks antiviral efficacy.

**Figure 1:**
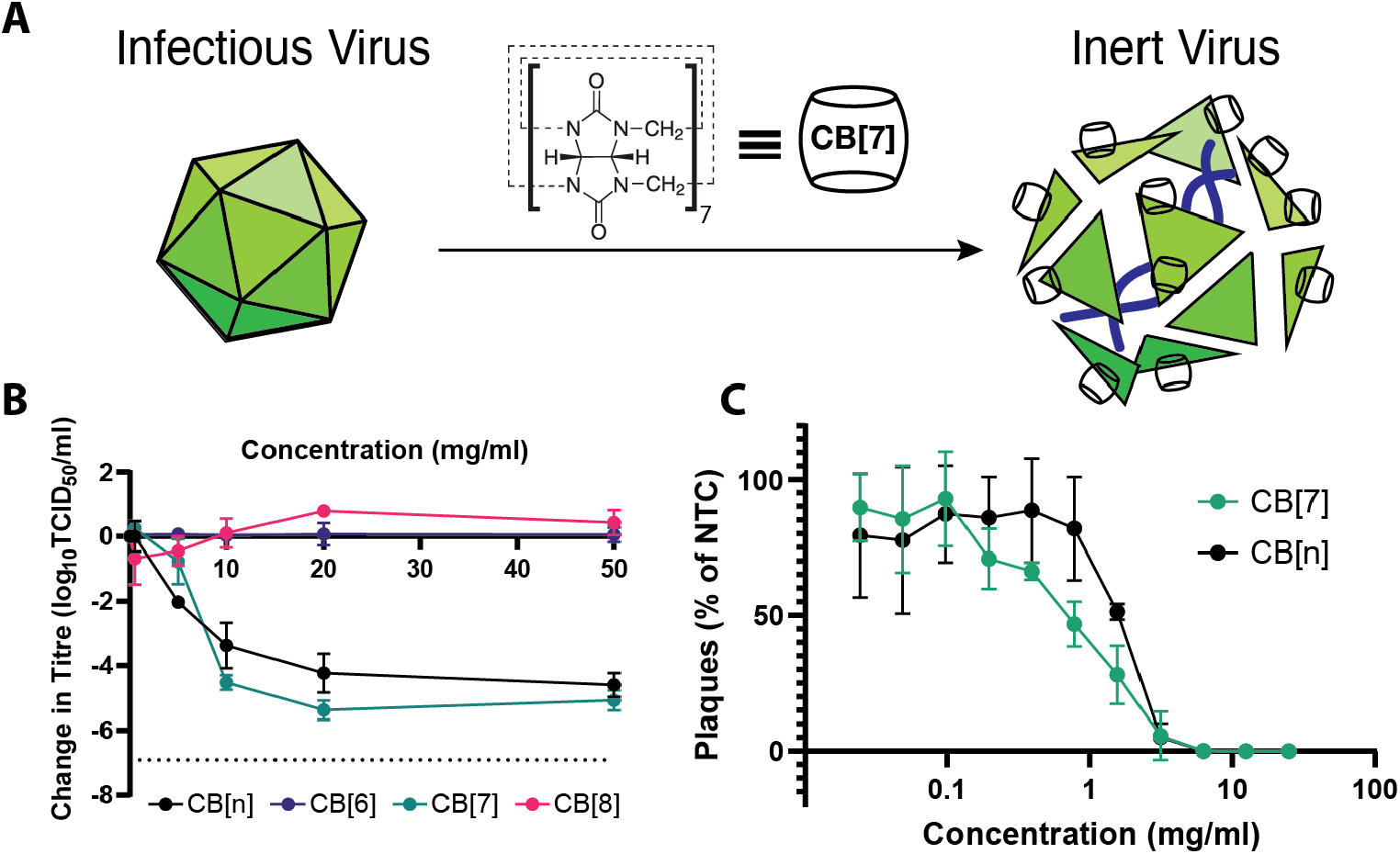
A) Graphic illustration depicting the virucidal antiviral effect of CB[7]. B) Data from TCID_50_ assays with various CBs against HSV-2, showing a decrease in HSV-2 titre when mixed with CB[n] and CB[7] but not with CB[6] and CB[8] (n=3). C) Dose-response assay against HSV-2 indicating that the IC50 value for CB[7] is lower than that for CB[n] (n=3). In all instances dotted lines indicate limits of detection, and error bars indicate standard deviation.

The effectiveness of CB[n] being lower than that of CB[7] is likely due to the fact that CB[n] contains a majority of the less effective CB[6] and CB[8] homologue.

Once the antiviral nature of CB[n] and CB[7] had been established, we used dose-response assays to quantify the effect. Such assays allow the half maximal inhibitory concentration (IC50) to be determined. Dose-response assays show that CB[n] has an IC50 of 1.5 mg/mL and CB[7] has an IC50 of 1.3 mg/mL (Figure 1C).

Additional experiments were performed via confocal fluorescence to confirm the effectiveness of CB[7] and CB[*n*] against HSV-2. This study showed that while virus was still detectable following 1:1 solution mixing with 0 and 0.5mg/ml CB[7], at higher levels (1mg/ml and above) no virus was subsequently detected following incubation on cells (Figure 2). In addition, when virus was mixed with CB[*n*], no virus was detected at 10mg/ml and above, although increasingly cellular damage, perhaps due to undissolved solid particles, is observed at 20mg/ml and particularly 50mg/ml (see ESI Figure S1). Although no viral presence was observed via confocal microscopy at 1mg/ml CB[7], TCID_50_ assays indicate some will still be present. This discrepancy can be explained through the fact that 5mg/ml CB[7] is associated with an approximately 0.8-log reduction, which may be enough to reduce the virus to a level that none was observed within the five blind-selected field-of-views of the monolayer.

**Figure 2:**
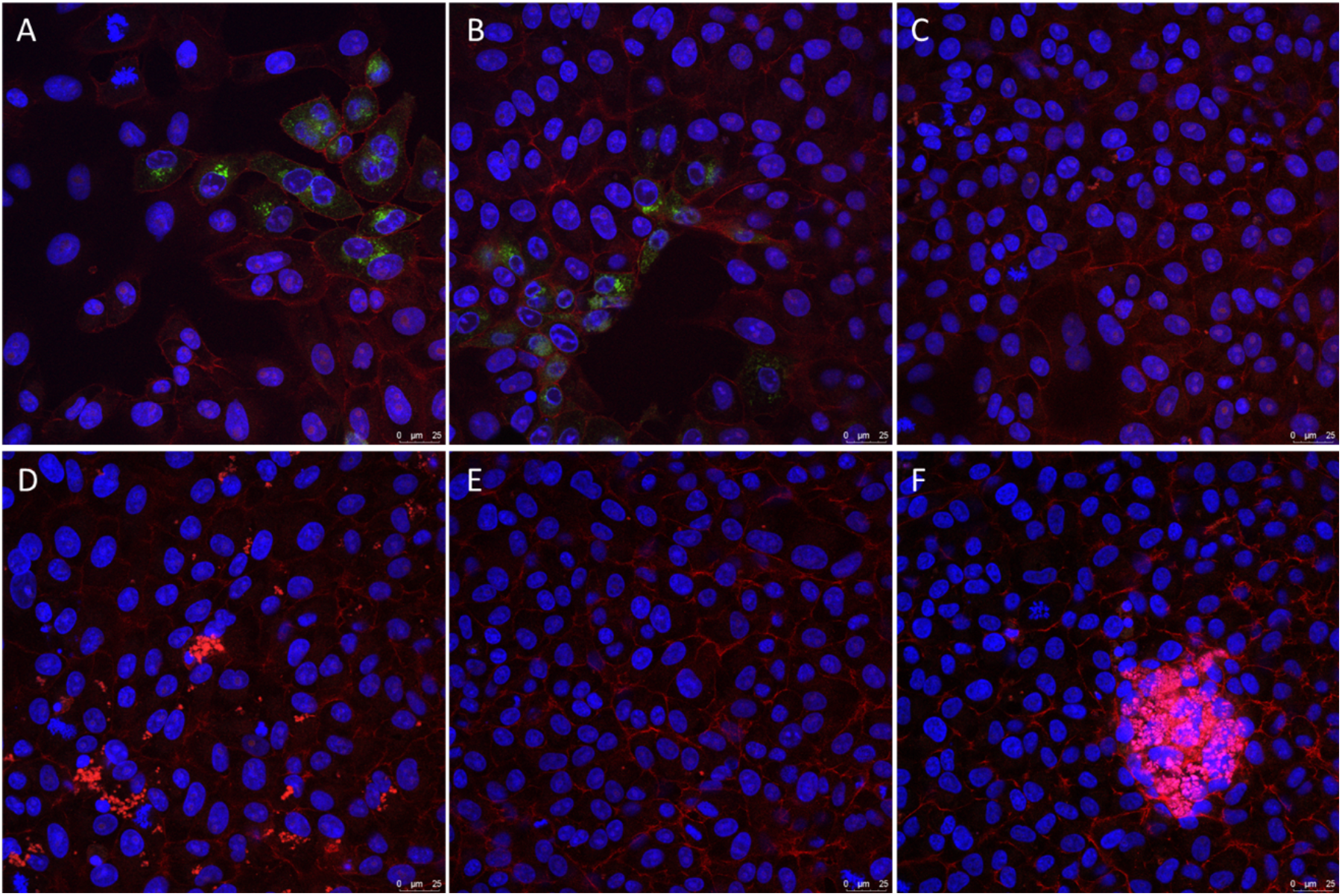
HSV-2 was mixed with different concentrations of CB[7] (A: 0mg/ml, B: 0.5mg/ml, C: 5mg/ml, D: 10mg/ml, E: 20mg/ml, F: 50mg/ml) and applied to cells, before incubating for 24 hours. HSV-2 in green, phalloidin in red, cell nuclei in blue.

### Mode of Action

In order to confirm that inhibition is due to supramolecular binding of the CB to the virus, we performed an additional assay, mixing CB[7] with 1-Adamantylamine (ADA). ADA has one of the strongest binding affinities (1.7 +/− 0.8 × 10^14^ M^−1^) [45] with CB[7], meaning the CB[7] cavity would be occupied and no binding to the virus could occur (Figure 3A). Different concentrations of 1:1 CB[7]:ADA were prepared and mixed with virus followed by TCID_50_ assays to determine viral titre. We observed that the 1:1 mix of CB[7]:ADA had no antiviral efficacy over the entire range (Figure 3B). Interestingly, ADA has antiviral efficacy of its own [46, 47]; however when mixed at equimolar ratio to CB[7] the antiviral efficacy of both compounds is neutralised. We performed a similar experiment with ferrocene, another strong binder with the CB[7] cavity (>10^6^ M^−1^) [48, 49] and one that lacks any antiviral properties. Complexation with ferrocene similarly eliminated the antiviral efficacy of the CB[7], reinforcing our hypothesis that the cavity of CB[7] is fundamental to its antiviral properties.

**Figure 3:**
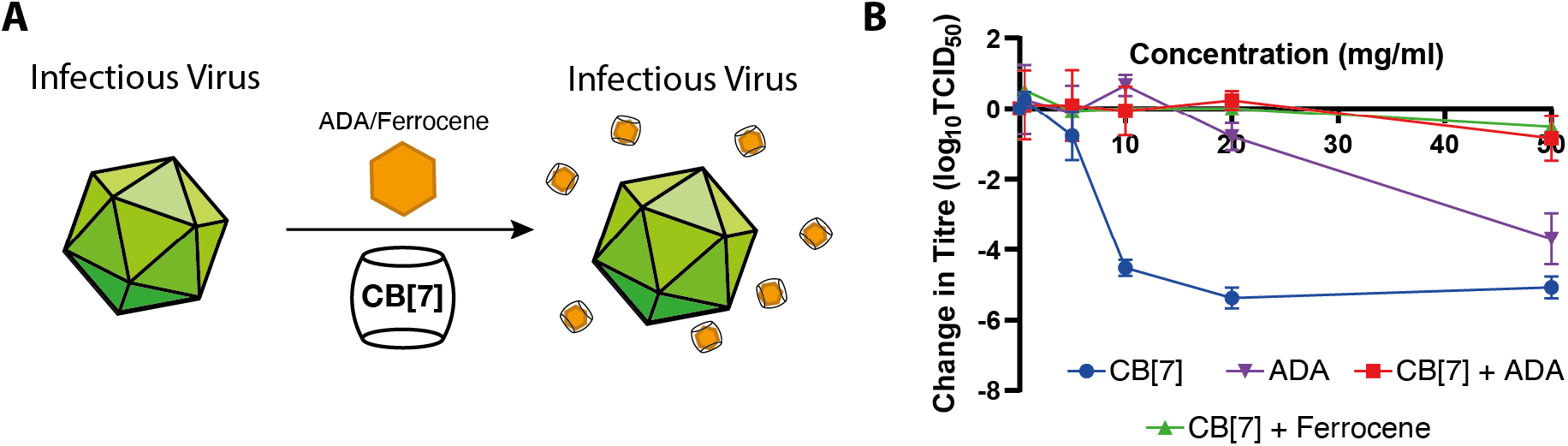
A) Graphic illustration depicting the ability of ADA or ferrocene to block the cavity of CB[7] and eliminate the antiviral effect. B) Data from TCID_50_ assays, illustrating the drop in titre when HSV-2 is exposed to CB[7] (n=3) or ADA alone (n=1), but no decrease in titre is observed when HSV-2 is mixed with CB[7] + ADA (n=1) or CB[7] + Ferrocene (n=1).

While previous assays confirm that CB[7] has antiviral properties, the exact mode of action was still unclear. CB[7] has been previously suggested to work by cellular internalization and binding to polyamines [36]. In order to further elucidate the antiviral mechanism, virucidal assays were performed with CB[7] against HSV-2. A virucidal assay is used to distinguish between destructive (virucidal) and non-destructive (virustatic) interactions between antivirals and viruses. In this instance, CB[7] was mixed with HSV-2 for 60 mins before being serially diluted across a 96-well plate. If the interactions between the CBs and virus were reversible non-destructive binding events, then serial dilution would cause removal of the CB from the viral surface and viral plaques would be observed at low dilutions. However, we observe that after serial dilution there is no recovery of virus, and a greater than 2-log reduction in viral titre, indicating that the binding of CB[7] to HSV-2 is indeed virucidal (Figure 4A). We also observe a virucidal mechanism with other viruses (*vide infra*).

**Figure 4:**
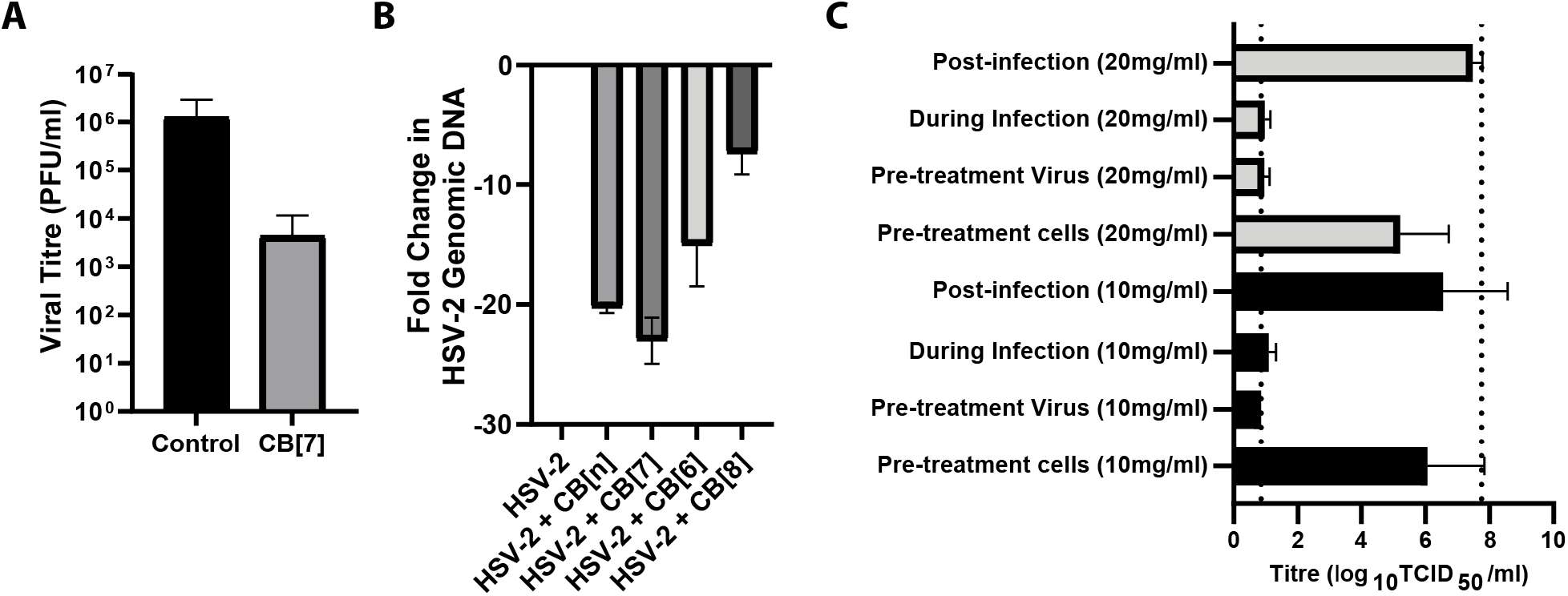
A) Virucidal assay data showing a 2-log reduction in viral titre with CB[7], indicating CB[7] acts virucidally (n=3). B) DNA-exposure assay shows that CB[7] and CB[n] are associated with release of DNA from the virus. C) Time of addition study suggests that CB[7] only inhibits HSV-2 titre when added simultaneously with virus to cells, and not when added before or after viral exposure (10 mg/ml in grey, 20 mg/ml in black) (n=3).

In order to confirm that the mode of action of the CB was extracellular, DNA exposure assays were performed. Such assays confirm the breakdown of the HSV-2 viral capsid and release of the genome in a cell free environment. This is critical to show that the mechanism is not via disruption of intracellular processes, such as viral replication pathways. The genome exposure assay includes a DNAse treatment step after CB[7] exposure – if the genome has been released (as would be the case for a virucide), it will be accessible to further enzymatic degradation thus unable to be amplified and detected via qPCR. Conversely, any viruses left intact by antiviral treatment will retain their genome, which can be detected via qPCR amplification. Data from this experiment shows that CB[n] and CB[7] were associated with significant drops in genome detection, suggesting that CB[7] does degrade the virion capsid and expose the genome to DNase treatment. Negative controls and CB[8] treatment did show detectable DNA, indicating that the viral capsid was not broken down (Figure 4B). CB[6] treatment was also associated with an unexpected drop in genomic DNA detection. Subsequent controls suggested that although a high enough CB concentration can affect qPCR results, not enough CB was present in any sample tested to change the Ct value (see ESI Figure S2A, B). This was investigated by adding various concentrations of CB[6], CB[7] and CB[8] to qPCR master mixes and observing the impact on Ct value (we showed higher concentrations of all CBs are associated with higher Ct values). In addition a positive control was performed, using Pseudomonas DNA in qPCR master mixes and adding the same volume of extracted viral DNA (that had initially been exposed to CB treatment). This had no effect on the Ct value. It is important to note that this assay illustrates that cucurbiturils can interact virucidally with viruses in a completely cell-free environment.

To elucidate further the antiviral mode of action of CBs, a series of time-of-addition studies was performed. We observed an antiviral effect when CB[7] was added directly to cells simultaneously with virus, or when virus stock was pre-treated with CB[7]. However, we observed no viral inhibition when the CB[7] was added to the cells and later removed (cell monolayers washed twice with PBS) prior to viral introduction. In addition, an assay was performed that allowed for infection followed by a period of CB exposure (to allow for potential CB-cell internalisation) before removal of the CB prior to incubation and viral replication/plaque formation. No antiviral effect was observed, further indicating that the CB binds directly to the virus in suspension (Figure 4C) with a virucidal mechanism. These assays strongly suggest that, in these systems at least, CB[7], is not internalised by the cells and thus does not disrupt the viral replication cycle, as previously reported [36].

### Broad Spectrum Efficacy

Having confirmed that CBs are effective antivirals against HSV-2, we also investigated if they were antiviral against other viruses. We began by testing CB[n] and CB[7] against respiratory syncytial virus (RSV). TCID50 assays (with a five-minute contact time between virus and antiviral, as previously) confirmed that both were also antiviral against RSV, with a 2-3 log reduction in viral titre at the highest concentrations. As previously, CB[6] and CB[8] were ineffective (Figure 5A), and the difference on average between CB[7] and CB[6]/CB[8] was statistically significant (adjusted p value <0.0001 for both). This was again quantified more precisely via dose-response assays, which showed that CB[7] has an IC50 value of 0.16 mg/ml and CB[n] also has an IC50 value of 0.16 mg/ml (Figure 5B). Virucidal assays were also performed with RSV and showed a 2-log reduction in viral titre (ESI Figure S3), indicating that CB[7] has a virucidal mode of action for both RSV and HSV-2. Cucurbiturils were also tested against cytomegalovirus (CMV) and seemed to have an antiviral effect when used in TCID50 assays; as previously CB[7] showed the most efficacy (Figure 5C). CB[7] titre reductions were significantly different from those with CB[8] (adjusted p-value of 0.0161) but otherwise there were no significant differences detected between cucurbiturils. In addition, both CB[7] and CB[n] seemed to be relatively ineffective when used in a dose-response assay, with IC50 values of 10.40mg/ml and 0.78mg/ml respectively (Figure 5D).

**Figure 5:**
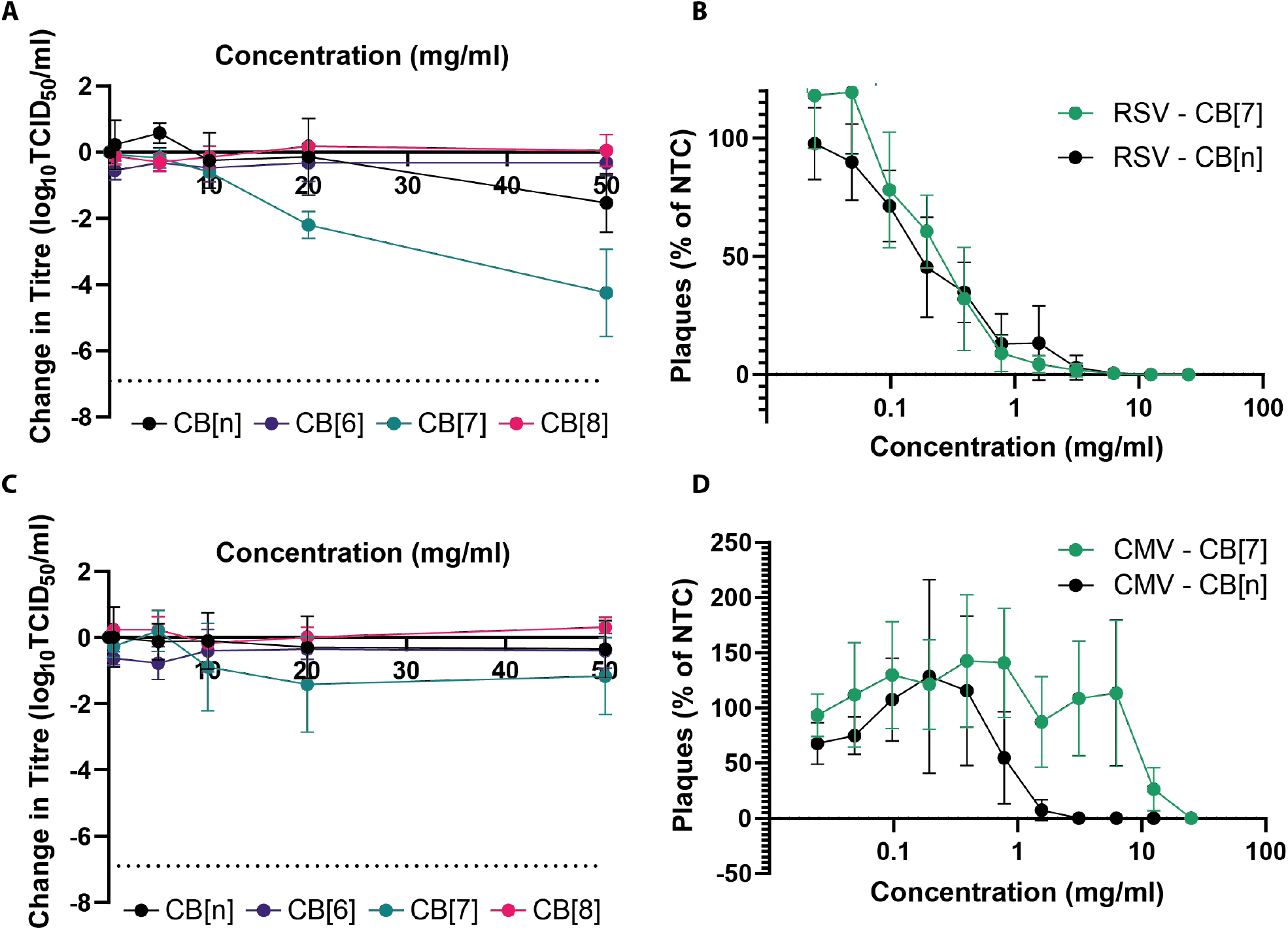
A) TCID_50_ assays against RSV showed that CB[7] and CB[n] are associated with large reductions in titre, particularly above 20mg/ml (n=3).B) Dose-response assays re-confirmed the effectivity of both CB[7] and CB[n], with both seemingly similarly effective when utilised in this assay (n=3). C) A limited reduction in titre was observed via TCID_50_ assay when CMV was mixed with different concentrations of CB[7]. CB[6], CB[8] and CB[n] appeared ineffective (n=3). D) Dose-response experiments with CMV showed that CB[7] (n=4) and CB[n] (n=4) display limited antiviral properties when tested in this manner.

Murine norovirus strain 1 (MNV1) (a surrogate for pathogenic human norovirus) was also investigated. Unlike previous viruses tested against, MNV1 is a non-enveloped virus. TCID50 assays demonstrated that no cucurbituril (including CB[7] and CB[n]) functions as an effective antiviral against this virus (ESI Figure S4). Subsequent statistical analysis found no significant difference between any cucurbituril homologue used in this experiment. Dose-response assays could not be performed for murine norovirus as it did not form plaques in the same manner as other viruses tested. Further experiments would be needed to determine why CB[n] proved ineffective against this virus, especially as efficacy was observed against a different MNV strain (MNV Berlin S99) (vide infra).

Human coronavirus are viruses of potential future pandemic concern. Therefore, we investigated the effects of CB[n] against two human coronaviruses; human coronavirus (strain OC43) and SARS-CoV-2 (Switzerland/GE9586/2020). OC43 was tested against CB[7] and CB[n] in a dose-response assay. Both CB[7] and CB[n] showed considerable efficacy against OC43, with IC50 values of 0.65mg/ml and 2.8mg/ml respectively (Figure 6A). Efficacy of cucurbiturils against SARS-CoV-2, the human coronavirus responsible for the COVID-19 pandemic, were also investigated. Titration assays against SARS-CoV-2 demonstrated that CB[n] and CB[7] are both antiviral against SARS-CoV-2, but to different extents. CB[n] has a more limited efficacy, even at a 50mg/ml concentration, whilst CB[7] is highly effective (approximately 6 log reduction) at 30 mg/ml and above. CB[8] has no antiviral effect (Figure 6B). These experiments suggest that CB[7] has broad efficacy against different human coronaviruses.

**Figure 6:**
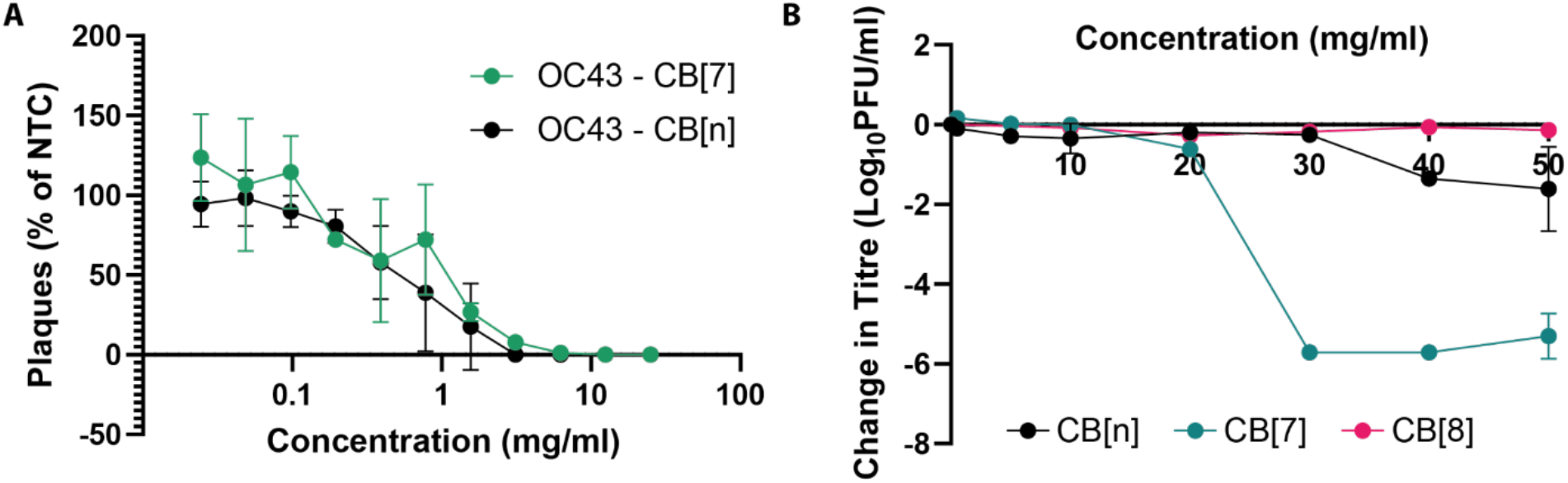
A) Dose-response assay performed against human coronavirus OC43. CB[7] (n=3) and CB[n] (n=3) appear similarly effective. B) CB[7] greatly reduces SARS-CoV-2 viral titre at 30mg/ml concentration and above (n=2). CB[n] displays a more limited antiviral effect over the same concentration range (n=2). CB[8] has no effect (n=1).

To simultaneously broaden the range of viruses tested against and further confirm the virucidal mode of action, a European standardized method was performed (EN 14476) in an accredited laboratory. EN 14476 specifically investigates products in suspension for use in disinfection in a medical context. Five viruses were investigated utilising this method; poliovirus type 1, murine norovirus, adenovirus type 5, modified vaccinia virus and feline coronavirus. A maximum of 5 mg/ml was used in this study to emulate concentrations used in CB-containing commercial products. The results from these experiments indicate that polio virus and adenovirus were not susceptible to CB[n], whereas feline coronavirus, vaccinia virus and murine norovirus were susceptible to CB[n] (all three showed an approximately 1 log reduction in titre at the highest concentrations tested) (Figure 7A). Polio virus, adenovirus and murine norovirus are all non-enveloped viruses; the presence or absence of an envelope is an important characteristic of a virus species and may be linked with the efficacy of CB[n]. Here we observe that while murine norovirus was inhibited by CB[n], polio virus and adenovirus were not.

**Figure 7:**
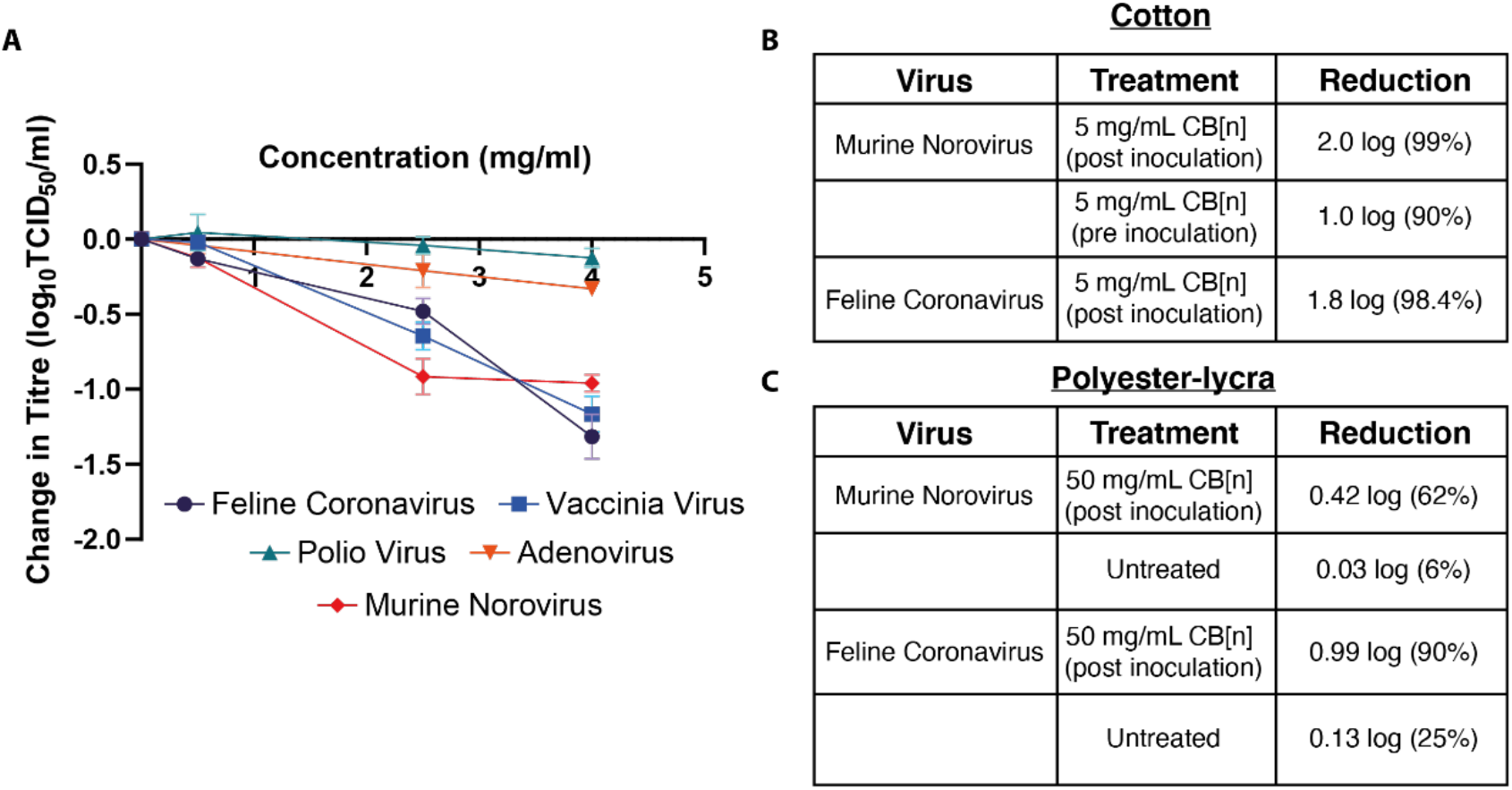
A) TCID_50_ assays conducted according to EN 14476 were performed after mixing viral suspensions with antiviral suspensions. Of all five viruses tested, only feline coronavirus, vaccinia virus and murine norovirus were inhibited by the CB[n] formulation (n=2). B) Surface testing of CB[n] formulations demonstrated some efficacy against murine norovirus and feline coronavirus. C) effect of pre-treated polyester-lycra on murine norovirus and feline coronavirus.

As we had confirmed that the CB mode of action was virucidal, similar to disinfectants such as bleach, this broadens the scope of potential applications. Unlike bleach and other virucides CBs are not damaging to surfaces such as textiles and have been demonstrated to be not irritating to skin or eye [50]. Therefore, to determine their possible function as a surface disinfectant, another standardised testing assay (ISO 18184) was used and performed at an accredited laboratory. Murine norovirus and feline coronavirus were again used as surrogates for human norovirus and human coronaviruses. Figure 7B shows that a formulation containing 5 mg/mL CB[*n*] leads to a 98.4% reduction in feline coronavirus when added onto cotton inoculated with the virus (post-inoculation) and a 99% reduction in murine norovirus (again post-inoculation). It was also observed, with murine norovirus, that a 90% reduction in viral titre was achieved when the textile was pre-treated with the CB[*n*] formulation (Figure 7B). In addition, further surface testing experiments were performed with treated fabric (polyester-lycra) at an accredited laboratory. These ISO 18184 surface tests again found a reduction in viral titre when the fabric was pre-treated with 5mg/ml CB[*n*]; feline coronavirus was reduced by 90% and murine norovirus by 62% (Figure 7C).

## Discussion

Through these results we have shown that cucurbit[n]urils are supramolecular broad-spectrum virucidal antivirals. Using HSV-2 as a model system we confirmed antiviral efficacy, dose effect, and a virucidal mechanism. Using guests with a strong binding affinity for the CB cavity we were able to show that this effect is on account of direct cavity binding to the virus. We were then able to broaden the range of viruses investigated to include several other species, including those of significant human-health concern, such as RSV and SARS-CoV-2, again confirming an antiviral effect. Using solution and surface (textile) European standardized testing methods, and surrogate viruses for some of the most dangerous human viruses, we further showed the significant potential of CBs as destroy-on-contact (virucidal) antivirals. This extracellular mode of action broadens the scope and real world uses of CB[n]s as antivirals, to potentially include topical treatments, prophylaxes, soft surface (textile) disinfection and through aerosolisation to deactivate airborne viruses.

## Materials and Methods

### Experimental Design

The study aimed to investigate whether cucurbiturils functioned as effective antiviral compounds, through the use of various cellular assays. Once efficacy was demonstrated, further experiments were conducted to establish the mechanism by which cucurbit[7]uril acts antivirals. This involved a range of cellular assays, but also a non-cellular assay (DNA exposure).

### Cucurbituril Stocks

Cucurbituril stocks (6, 7, 8 and a mixture of all three) were provided courtesy of Aqdot Ltd in dry powdered form. They were mixed with sterile deionized water (dH2O) to form an initial 5% stock concentration (0.05g of powder/ml dH2O, a 50mg/ml solution). These were then diluted appropriately in sterile dH2O to form additional 2%, 1%, 0.5% and 0.05% stocks. dH2O was sterilized via filtration with a disposable syringe filter (Merck Millipore Ltd, Cork, Ireland) before use.

### Viruses

Herpes Simplex Virus, serotype two (HSV-2) and respiratory syncytial virus (RSV) samples were originally isolated, verified and kindly donated by the University of Manchester School of Medical Sciences (Dr Carol Yates). Additional stocks were grown on Vero cells in-lab and stored at −80°C. Murine norovirus (strain 1) was purchased from ATCC before stocks were grown up on RAW 264.7 cells in-lab.

Coronavirus (strain OC43) was originally isolated, verified and kindly donated by University of Manchester School of Biological Sciences (Prof Pamela Vallely and Prof Paul Klapper). Additional stocks were grown on MK1-Lu cells in-lab and stored at −80°C. SARS-CoV-2 (strain SARS-CoV2/Switzerland/GE9586/2020) was obtained from a clinical specimen in the University Hospital in Geneva using Vero-E6 cells and passaged twice before use in experiments. For experiments, virus was propagated in Vero C1008 (clone E6) cells. Virus was handled appropriately in CL-3 laboratories.

### Cell Culture

All cell culture was performed using aseptic techniques in a class II microbiological safety cabinet (MBSC-II). All tissue culture media was supplemented with 1% penicillin/streptomycin (P/S) (Merck Life Science UK Ltd, Dorset, United Kingdom) and 10% heat inactivated foetal calf serum (FCS) (Merck Life Science UK) unless stated otherwise.

Vero cells were kindly donated by the University of Manchester School of Medical Sciences (Dr Carol Yates). They were maintained in Dulbecco’s Modified Eagle Medium (DMEM) modified with High Glucose, L-Glutamine, Phenol Red and Sodium Pyruvate (Thermo Fisher Scientific, Loughborough, UK). Cells were cultivated at 37°C, 5% CO_2_ in 75cm^2^ flasks and passaged in a 1:6 ratio when confluent. Vero C1008 (clone E6) cells, used for SARS-CoV-2 studies, were grown in DMEM modified with high glucose and Glutamax.

RAW 264.7 cells (Merck Life Science) were maintained in DMEM modified with phenol red and L-glutamine. Cells were cultivated at 37°C, 5% CO_2_ in 75cm^2^ flasks and passaged via cell scraping when confluent.

Mv-1 Lu cells (mink lung epithelial cells) were kindly donated by University of Manchester School of Biological Sciences (Dr Pamela Vallely) and were maintained in DMEM modified with high glucose, sodium pyruvate, phenol red and L-glutamine. Cells were cultivated at 37°C, 5% CO_2_ in 75cm^2^ flasks and passaged in a 1:6 ratio when confluent.

### Titration by TCID_50_ Assay

Samples were titrated in 96-well flat bottom plates (Thermo Fisher Scientific), in quadruplicate. The first four wells of the first column were filled with 180μl of appropriate media (see Cell Culture) and all other wells were filled with 100μl of media containing appropriate cells at 2×10^5^/ml concentration. After cells adhered to the plate, wells A1-D1 then had 20μl of sample added, before the contents were mixed via pipetting; 100μl of mixed sample was then transferred to the next four wells and mixed again. This serial dilution was repeated across the plate, with the exception of wells A12-D12, which were left unexposed to sample as a negative control. Plates were then incubated for an appropriate length of time depending on the virus being investigated. Virus titre was determined by cytopathic effects observed in wells via light microscopy examination. Titre was calculated using the Spearman and Käber method.

### Guest-host Chemistry Assay

CB[7] powder stock was mixed with adamantylamine in a 1:1 molar ratio and resuspended at 50mg/ml in sterile dH2O. Samples from this solution were then diluted further to form stocks of 50mg/ml, 20mg/ml, 10mg/ml, 5mg/ml and 0.5mg/ml.

### Dose-Response Assays

Cells were seeded at 170,000/ml, with 500μl of cell and media mixture added to each well of a 24 well plate (Corning, NY, USA). They were incubated at 37°C overnight or until cells were 90-100% confluent. Sterile Eppendorf tubes were then prepared containing 570μl of antiviral material mixed with DMEM to provide the desired concentration of antiviral. A 1:100 dilution of virus stock was also prepared; 30μl of this diluted virus stock was added to each Eppendorf containing 570μl of antiviral/media mix. An addition Eppendorf tube was also prepared containing just 570μl DMEM plus 30μl diluted virus stock to act at the non-treatment control (NTC). Eppendorfs containing antiviral-virus mix and NTC Eppendorf were incubated for one hour at 37°C. Then all media was removed from the 24-well plate and 200μl of each antiviral-virus mix added (in duplicate). The plate was then incubated for one hour at 37°C. Finally, the antiviral-virus mixes were removed by pipetting and 500μl of methylcellulose DMEM was added to each well. The plate was then incubated under the formation of visible plaques.

### Virucidal Assays

Cells appropriate to the virus were seeded into a 96-well plate at 15,000 cells per well and left until 90-100% confluent. Virus stock (55μl) was incubated with the desired amount of antiviral agent to achieve the IC90 concentration in 55μl total volume (made up with phosphate buffered saline (PBS; Cambridge Bioscience, United Kingdom)). A non-treatment control was also created, with 55μl virus stock mixed with 55μl PBS. Both virus plus antiviral and non-treatment controls were incubated for 1 hour at 37°C. Then six 2ml Eppendorf tubes were prepared, each containing 450μl of appropriate cell culture medium (three were used with the antiviral, three with the non-treatment control). After incubation, 50μl was taken from the virus plus antiviral mix and added into the first Eppendorf tube before being resuspended 4-5 times. 50μl was then serially diluted into the other two tubes, leaving three Eppendorfs at 1:20, 1:200 and 1:2000 dilution respectively. This process was also repeated for the non-treatment control. 50μl from the virus plus antiviral mix (1:20 dilution) was then added to wells A1 and A2 of the 96-well plate. 50μl from the 1:200 virus-antiviral mix was added to wells A3 and A4, and 50μl from the 1:2000 virus-antiviral mix was added to wells A5 and A6. 50μl from these wells was then serially diluted down the plate to row G. In well H1, 50μl of the originally incubated virus plus antiviral stock was added, resuspended, and serially diluted to well H6. This process was repeated for the non-treatment control in wells A7-A12. The plate was then incubated for one hour at 37°C. All medium was then discarded and replated with 100μl of methylcellulose DMEM. The plate was then incubated at 37°C until visible plaques were formed, before being stained with crystal violet to allow plaque-counting.

### Genome Exposure Assay

#### Enzymatic Treatment

For HSV-2, 100μl of viral sample was combined with 100μl of antiviral sample in a sterile Eppendorf tube. A positive control was also included where 100μl of virus stock is mixed with 100μl of sterile deionized water. All samples were then incubated for five minutes at 37°C in 5% CO_2_. Following co-incubation of virus and antiviral, DNA concentration of each sample were measured using a NanoDrop Lite Spectrophotomer (ThermoFisher Scientific). The DNA or RNA concentration of each sample was noted. For DNA-genome viruses, 20μl of 10X TURBO™ Dnase Buffer was added to each sample. In the sample with the highest amount of DNA (as measured by the spectrophotometer), 1μl of TURBO™ Dnase was added per 1 μg DNA present. An equal volume of Turbo Dnase was also added to all other samples in the batch. Samples were then incubated for 30 minutes at 37°C in 5% CO2. Finally, EDTA (Merck Life Sciences) was added to each sample at a final concentration of 15 mM, and heated at 75°C for 10 minutes to deactivate the Turbo Dnase.

#### Genome Extraction

Extraction was performed using a PureLink^®^ Viral RNA/DNA Mini Kit (Invitrogen) according to manufacturer’s instruction. Briefly, 25μL Proteinase K was added, before adding 200μL Viral Lysis Buffer. Samples were briefly vortexed and incubated at 56°C for 15 minutes. 250μL molecular grade ethanol (Merck Life Sciences) was then added to each sample before brief vortexing and incubation for 5 minutes at room temperature. Samples were then added to a Viral Spin Column in a collection tube before being centrifuged in a benchtop microcentrifuge at 6800 × g for 1 minute (all subsequent centrifugation steps assume this speed unless otherwise specified). Flow-through was discarded and 500μL Wash Buffer (WII) was added, before being centrifuged for 1 minute. Flow-through was discarded and another 500μl of Wash Buffer added to each sample, before being centrifuged for one minute. The spin column was then centrifuged in a clean collection tube for 1 minute to remove any residual Wash Buffer (WII). Each spin column was then placed in a collection tube and genetic material eluted with 50μL sterile Rnase-free water. Samples were then incubated at room temperature for 1 minute before being centrifuged for 1 minute to elute nucleic acids. Purified viral DNA was stored at –80°C before being amplified in qPCR.

#### qPCR Amplification

qPCR was performed with a PowerUp™ SYBR Green™ Master Mix (Applied BioSystems). See Table 1 for primers used for qPCR; all primers used were purchased from Eurofins Europe. Samples for qPCR were made up to 20μl total before being analysed with the qPCR machine (Applied Biosystems StepOne Plus™, ThermoFisher Scientific) and associated software. The PCR sequence used was as follows: holding stage (95°C) for ten minutes, followed by 40 cycles of 95°C for fifteen seconds, and 60°C for one minute. This was followed by a single melt-curve stage of 95°C for 15 seconds and 60°C for one minute.

**Table 1:**
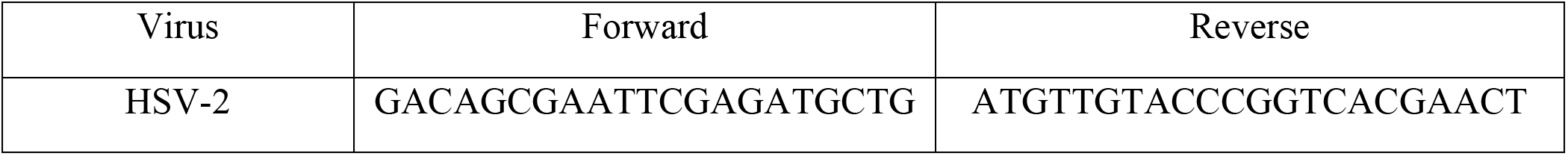
List of Primers used in qPCR Experiment

#### Bacterial qPCR samples

Experiments were conducted using Pseudomonas aeruginosa (AP) strain ATCC PA01. P. aeruginosa was incubated overnight on nutrient agar plates. Bacterial DNA was extracted using QIAamp DNA mini kit (Qiagen), using one bacterial colony. Primers for amplification were; forward primer (27F), reverse primer (1492R) (Eurofins Genomics).

### Time of Addition Study

Materials (10μg or 20μg) were added on cells 1 hour before infection, during infection or after infection, with viruses added using an MOI of 0.01, using a method described previously [19, 51]. Viral titres were then quantified by TCID_50_ assay.

### Immunofluorescence Experiment

13mm-diameter glass coverslips (Fisher Scientific) were added to 24-well plates (Corning Costar, Fisher Scientific). Cells of interest were seeded at a density of 2×10^5^/ml and left to adhere overnight at 37°C, 5% CO_2_. Virus stock of known titre was mixed with antiviral samples and added to cells for sixty minutes to allow for adsorption. After this, infectious medium was removed and the well washed with 1ml PBS applied by pipette, before 1ml fresh tissue culture medium was added. After the required incubation period (24 hours for HSV-2) tissue culture medium was removed and the infected cells were fixed in 10% formalin for 1 hour. After fixation, coverslips were transferred to a new 24 well plate and stored in PBS at 4°C until labelling was performed. Labelling began with the removal of PBS and addition of 0.1% Triton X-100 (Merck Life Sciences) for approximately 15 minutes. After this, coverslips were washed once with PBS, before non-specific binding was blocked by the addition of 0.5% PBS/Bovine Serum Albumin (BSA) for 30 minutes. 0.5% PBS/BSA was then removed and primary antibody (anti-HSV-2 antibody 2C10, Abcam, United Kingdom), diluted in 0.5% PBS/BSA, was added for 60 minutes. After this, coverslips were washed three times in PBS, before incubation with secondary antibody (Alexa Fluor™ 488, ThermoFisher Scientific) (diluted in 0.5% PBS/BSA) for 60 minutes. Next, coverslips were again washed three times in PBS, before being incubated with Alexa Fluor™ 568 Phalloidin (1:60 dilution) for 20 minutes. Phalloidin was then removed and coverslips were incubated for ten minutes in 1:10,000 DAPI (Merck Life Sciences) solution (made up in deionized water). A drop of VECTASHIELD (Vectorlabs) mounting medium was then applied to a glass slide; coverslips were dipped in distilled water with fine forceps, and then dried by being placed vertically on a paper towel. The coverslip was then placed, cells down, onto the mounting medium droplet. Coverslips were sealed with application of CoverGrip™ Sealant (Cambridge Bioscience) around the circumference, before being allowed to dry. Once dry, samples were stored in a dark at 4°C before being viewed on a Leica confocal microscope, with LAS X software (Leica Microsystems).

### Solution Testing (EN 14476)

A sample of the supplied test product (CB[n] commercial formulation Oderase) was diluted in distilled water. This was added to a test suspension of viruses in a solution of interfering substance (final concentration 0.3 g/L bovine albumin). The mixture was maintained at 20°C ± 1°C and at different contact times (5 minutes and 1 hour, data combined above). At the end of this contact time, an aliquot was taken and the virucidal action in this portion was immediately suppressed by a validated method (dilutions of the sample in ice-cold cell maintenance medium). The dilutions were transferred into cell culture units either using monolayer or cell suspension. Infectivity tests were done either by plaque test or quantal tests. After incubation, the titres of infectivity were calculated according to Spearman and Käber or by plaque counting. Reduction of virus infectivity was calculated from differences of log virus titres before (virus control) and after treatment with the supplied product. The spectrum of test organisms was poliovirus (Type 1KSc), adenovirus (Adenoid 75), murine norovirus (strain S99 Berlin), vaccinia virus (ATCC VR-5) and feline coronavirus (Munich strain).

### Textile Testing (ISO 18184)

A 20mm × 20mm sample of test material was cut (overall mass should be 0.40g and was made up with extra layers of material as required). 9 control pieces were required and 6 test pieces. 3 pieces of each material were used to test the effect of the fabric on cells without virus (cytotoxicity), 3 control pieces were used to recover the starting titre of virus. The remaining pieces were inoculated with 200μl of virus at a concentration of ~10^7^ TCID_50_ (giving a final concentration of 10^5^) and left for the contact time of two hours. Following the contact time, the fabric was recovered in 20ml of cell culture media and enumerated onto an appropriate cell line. TCID_50_ was calculated following the appropriate incubation time. Antiviral activity was calculated by comparison of the antiviral test material to the immediate recovery from the control fabric.

Additional textile testing was carried out on polyester-lycra samples. In these instances, impregnated AqFresh textile was prepared by immersing samples of polyester-lycra in a 5 wt% AqFresh suspension in water (1L in 3L beaker) for 10 minutes while mixing at 300 rpm using a magnetic stirrer. Wet textile samples were recovered, then squeezed between two rollers to remove excess solution until no further solution could be removed. Textile samples were then dried in a convection oven at 110°C for 15 minutes. The AqFresh loading of the dried treated textile was determined gravimetrically to be 5 wt% on average. Control textiles were immersed in water and treated by the same procedure. Antiviral tests were conducted following ISO 18184.

### Statistical Analysis

Statistical analysis (for generation of IC50 values) was performed using GraphPad Prism software, version 9.1.1, using a non-linear fit of [Inhibitor] vs. response – variable slope (four parameters) in Graphpad Prism software. Furthermore, different cucurbituril treatments in the TCID_50_ assays were analyzed for statistical differences using the same version of GraphPad Prism software, using a 2-way ANOVA analysis with Tukey’s multiple comparisons.

## Acknowledgments

The authors would like to thank Francesco Stellacci at the Supramolecular NanoMaterials and Interfaces Laboratory in Switzerland for his help with all the SARS-CoV-2 related studies. The authors would also like to thank Prof. Pamela Vallely and Prof. Paul Klapper for their help with supply of cell lines and viruses. The authors would also like to thank Dr. Lee Fielding and Elisabeth Trinh for generously providing bacterial DNA and associated primers. Finally, the authors would also like to thank Dr. Nico Esselin for preparing the AqFresh textiles.

## Author contributions

LMJ, ES and LB performed experiments for Figures 1–6A. MG performed SARS-CoV-2 experiments for Figure 6B. BC, AH, RC commissioned testing for Figure 7. LMJ and STJ wrote the manuscript.

## Data and materials availability

All data needed to evaluate the conclusions in the paper are present in the paper and/or the Supplementary Materials. Additional data related to this paper may be requested from the authors.

## References

1. Smith, K.F., et al., Global rise in human infectious disease outbreaks. Journal of the Royal Society Interface, 2014. 11(101): p. 20140950.

2. Nandy, A. and S. Basak, Viral epidemics and vaccine preparedness. J Mol Pathol Epidemiol, 2017. 2: p. S1.

3. Bloom, D.E., D. Cadarette, and J. Sevilla, Epidemics and economics. Finance & Development, 2018. 55(002).

4. Rasul, I. The economics of viral outbreaks. in AEA Papers and Proceedings. 2020.

5. Bhella, D., The role of cellular adhesion molecules in virus attachment and entry. Philosophical Transactions of the Royal Society B: Biological Sciences, 2015. 370(1661): p. 20140035.

6. Lin, Q., et al., Sanitizing agents for virus inactivation and disinfection. View, 2020. 1(2): p. e16.

7. Baker, N., et al., Repurposing quaternary ammonium compounds as potential treatments for COVID-19. Pharmaceutical research, 2020. 37: p. 1–4.

8. Sokolova, A.S., et al., New quaternary ammonium camphor derivatives and their antiviral activity, genotoxic effects and cytotoxicity. Bioorganic & medicinal chemistry, 2013. 21(21): p. 6690–6698.

9. El-Sheekh, M.M., et al., Antiviral activity of algae biosynthesized silver and gold nanoparticles against Herps Simplex (HSV-1) virus in vitro using cell-line culture technique. International Journal of Environmental Health Research, 2020: p. 1–12.

10. Paul, A.M., et al., Delivery of antiviral small interfering RNA with gold nanoparticles inhibits dengue virus infection in vitro. The Journal of general virology, 2014. 95(Pt 8): p. 1712.

11. Shionoiri, N., et al., Investigation of the antiviral properties of copper iodide nanoparticles against feline calicivirus. Journal of bioscience and bioengineering, 2012. 113(5): p. 580–586.

12. Rosa Borges, A. and C.-L. Schengrund, Dendrimers and antivirals: a review. Current Drug Targets-Infectious Disorders, 2005. 5(3): p. 247–254.

13. Vacas-Córdoba, E., et al., Antiviral mechanism of polyanionic carbosilane dendrimers against HIV-1. International journal of nanomedicine, 2016. 11: p. 1281.

14. Vaillant, A., Nucleic acid polymers: broad spectrum antiviral activity, antiviral mechanisms and optimization for the treatment of hepatitis B and hepatitis D infection. Antiviral research, 2016. 133: p. 32–40.

15. Bai, H., et al., Macrocyclic compounds for drug and gene delivery in immune-modulating therapy. International journal of molecular sciences, 2019. 20(9): p. 2097.

16. Jazkewitsch, O., et al., Cyclodextrin-modified polyesters from lactones and from bacteria: an approach to new drug carrier systems. Macromolecules, 2011. 44(6): p. 1365–1371.

17. Singh, M., R. Sharma, and U. Banerjee, Biotechnological applications of cyclodextrins. Biotechnology advances, 2002. 20(5-6): p. 341–359.

18. Zabirov, N., et al., Antiviral activity of macrocyclic polyethers and their complexes with the alkaline metal salts of N-phosphorylated amides and thioamides. Pharmaceutical chemistry journal, 1991. 25(5): p. 321–324.

19. Jones, S.T., et al., Modified cyclodextrins as broad-spectrum antivirals. Science advances, 2020. 6(5): p. eaax9318.

20. Graham, D.R., et al., Cholesterol depletion of human immunodeficiency virus type 1 and simian immunodeficiency virus with β-cyclodextrin inactivates and permeabilizes the virions: evidence for virion-associated lipid rafts. Journal of virology, 2003. 77(15): p. 8237–8248.

21. Kim, K., Cucurbiturils and related macrocycles. Vol. 28. 2019: Royal Society of Chemistry.

22. Day, A., et al., Controlling factors in the synthesis of cucurbituril and its homologues. The Journal of organic chemistry, 2001. 66(24): p. 8094–8100.

23. Freeman, W., W. Mock, and N. Shih, Cucurbituril. Journal of the American Chemical Society, 1981. 103(24): p. 7367–7368.

24. Kim, J., et al., New cucurbituril homologues: syntheses, isolation, characterization, and X-ray crystal structures of cucurbit [n] uril (n= 5, 7, and 8). Journal of the American Chemical Society, 2000. 122(3): p. 540–541.

25. Nau, W.M., M. Florea, and K.I. Assaf, Deep inside cucurbiturils: physical properties and volumes of their inner cavity determine the hydrophobic driving force for host–guest complexation. Israel Journal of Chemistry, 2011. 51(5-6): p. 559–577.

26. Gong, W., et al., From Packed “Sandwich” to “Russian Doll”: Assembly by Charge-Transfer Interactions in Cucurbit [10] uril. Chemistry–A European Journal, 2016. 22(49): p. 17612–17618.

27. Kim, H.J., et al., Selective inclusion of a hetero-guest pair in a molecular host: formation of stable charge-transfer complexes in cucurbit [8] uril. Angewandte Chemie International Edition, 2001. 40(8): p. 1526–1529.

28. Walker, S., et al., The potential of cucurbit [n] urils in drug delivery. Israel Journal of Chemistry, 2011. 51(5-6): p. 616–624.

29. Saleh, N.i., A.L. Koner, and W.M. Nau, Activation and Stabilization of Drugs by Supramolecular pKa Shifts: Drugs-Delivery Applications Tailored for Cucurbiturils. Angewandte Chemie, 2008. 120(29): p. 5478–5481.

30. Hou, C., et al., Cucurbituril as a versatile tool to tune the functions of proteins. Israel Journal of Chemistry, 2018. 58(3-4): p. 286–295.

31. Khaligh, A. and D. Tuncel, Cucurbituril-assisted Supramolecular Polymeric Hydrogels, in Cucurbituril-based Functional Materials. 2019. p. 120–148.

32. Rana, V.K., et al., Cucurbit [8] uril-derived graphene hydrogels. ACS Macro Letters, 2019. 8(12): p. 1629–1634.

33. Buschmann, H.-J., K. Jansen, and E. Schollmeyer, The formation of cucurbituril complexes with amino acids and amino alcohols in aqueous formic acid studied by calorimetric titrations. Thermochimica acta, 1998. 317(1): p. 95–98.

34. Zebaze Ndendjio, S.A. and L. Isaacs, Molecular recognition properties of acyclic cucurbiturils toward amino acids, peptides, and a protein. Supramolecular Chemistry, 2019. 31(7): p. 432–441.

35. Chinai, J.M., et al., Molecular recognition of insulin by a synthetic receptor. Journal of the American Chemical Society, 2011. 133(23): p. 8810–8813.

36. Quan, J., et al., Cucurbit [7] uril as a Broad-Spectrum Antiviral Agent against Diverse RNA Viruses. Virologica Sinica, 2021: p. 1–12.

37. Ahmadi, V., et al., One-pot gram-scale synthesis of virucidal heparin-mimicking polymers as HSV-1 inhibitors. Chemical Communications, 2021. 57(90): p. 11948–11951.

38. Cagno, V., et al., Broad-spectrum non-toxic antiviral nanoparticles with a virucidal inhibition mechanism. Nature materials, 2018. 17(2): p. 195–203.

39. Minoshima, M., et al., Comparison of the antiviral effect of solid-state copper and silver compounds. Journal of Hazardous Materials, 2016. 312: p. 1–7.

40. Rakowska, P.D., et al., Antiviral surfaces and coatings and their mechanisms of action. Communications Materials, 2021. 2(1): p. 1–19.

41. Lee, J., S, Samal, N. Selvapalam, H.-J. Kim, and K. Kim. Acc. Chem. Res, 2003. 36: p. 621.

42. Masson, E., et al., Cucurbituril chemistry: a tale of supramolecular success. Rsc Advances, 2012. 2(4): p. 1213–1247.

43. Pan, S., S. Mandal, and P.K. Chattaraj, Cucurbit [6] uril: a possible host for noble gas atoms. The Journal of Physical Chemistry B, 2015. 119(34): p. 10962–10974.

44. Rajgariah, P. and A.R. Urbach, Scope of amino acid recognition by cucurbit [8] uril. Journal of Inclusion Phenomena and Macrocyclic Chemistry, 2008. 62(3): p. 251–254.

45. Moghaddam, S., et al., New ultrahigh affinity host− guest complexes of cucurbit [7] uril with bicyclo [2.2. 2] octane and adamantane guests: Thermodynamic analysis and evaluation of m2 affinity calculations. Journal of the American Chemical Society, 2011. 133(10): p. 3570–3581.

46. Danilenko, G., et al., Synthesis and biological activity of adamantane derivatives. IV. Viral inhibiting activity of some adamantylamines. Pharmaceutical Chemistry Journal, 1976. 10(6): p. 737–741.

47. Yang, J., et al., Synthesis and antiviral activities of novel gossypol derivatives. Bioorganic & medicinal chemistry letters, 2012. 22(3): p. 1415–1420.

48. Jeon, W.S., et al., Complexation of ferrocene derivatives by the cucurbit [7] uril host: a comparative study of the cucurbituril and cyclodextrin host families. Journal of the American Chemical Society, 2005. 127(37): p. 12984–12989.

49. Ong, W. and A.E. Kaifer, Unusual electrochemical properties of the inclusion complexes of ferrocenium and cobaltocenium with cucurbit [7] uril. Organometallics, 2003. 22(21): p. 4181–4183.

50. Toxicological Information. 2015 [cited 2021 14/12]; Available from: https://echa.europa.eu/registration-dossier/-/registered-dossier/29689/7/4/1.

51. Aoki-Utsubo, C., M. Chen, and H. Hotta, Time-of-addition and temperature-shift assays to determine particular step (s) in the viral life cycle that is blocked by antiviral substance (s). Bio-protocol, 2018. 8: p. e2830.

