## Supplementary Material for "Broad-spectrum extracellular antiviral properties of Cucurbit[n]urils"

### Confocal Immunofluorescence Microscopy images of Vero cell monolayers after 24hr exposure to CB[*n*] mixed with HSV-2

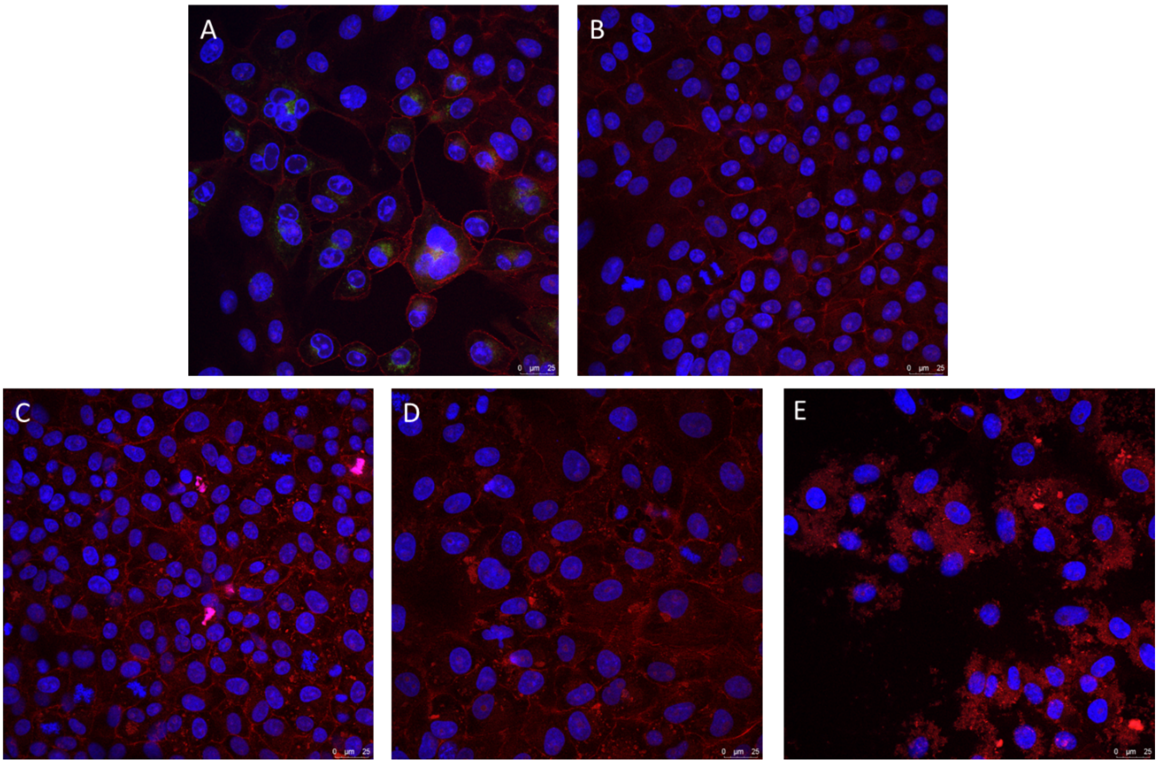


Figure S1: HSV-2 was mixed with different concentrations of CB[*n*] (A: 0mg/ml, B: 5mg/ml, C: 10mg/ml, D: 20mg/ml, E: 50mg/ml) and applied to cells, before incubating for 24 hours. Visual indications are that CB[n] has an antiviral effect at 5mg/ml concentrations and above. However, at 20mg/ml and particularly 50mg/ml, cellular damage becomes increasingly visible, with the monolayer broken apart at the highest concentrations (E). Cell nuclei in blue, phalloidin in red, HSV-2 in green.

### qPCR Controls to determine the effect of CBs on Ct values

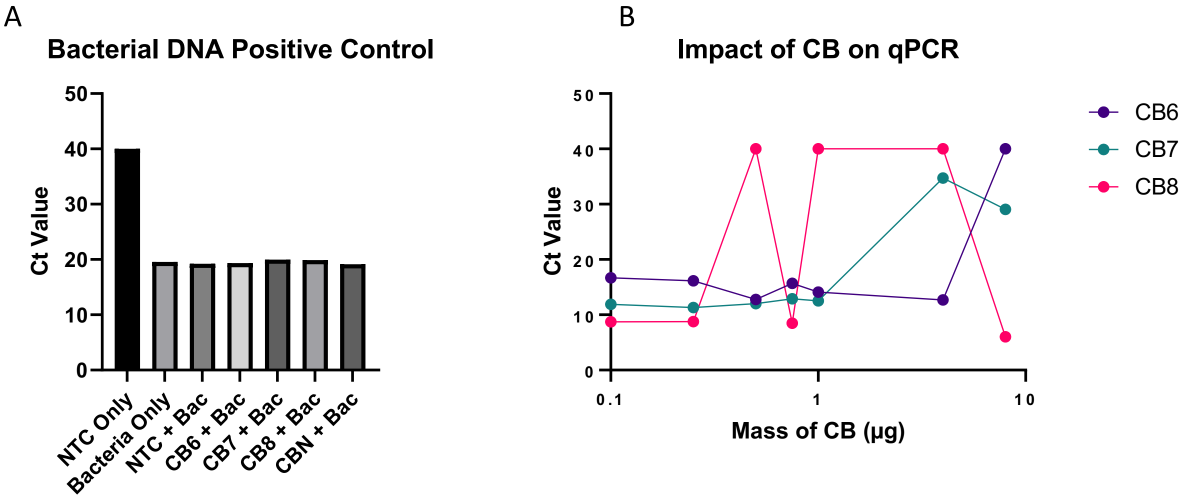


Figure S2: A) Bacterial DNA and primers were run as a positive control for qPCR. PCR primers did not amplify extracted HSV-2 viral DNA (left most bar). On all other occasions, extracted viral DNA samples, that had initially been mixed 1:1 with cucurbiturils, did not contain enough cucurbituril to artificially alter the Ct value. B) Interestingly, when cucurbiturils are added directly to the PCR mastermix, they can alter the Ct value, though only above certain concentrations (which vary depending on the homologue). For this experiment CBs were added directly to PCR master mixes containing extracted HSV-2 DNA (that had not been exposed to any cucurbituril treatment).

### Virucidal RSV data

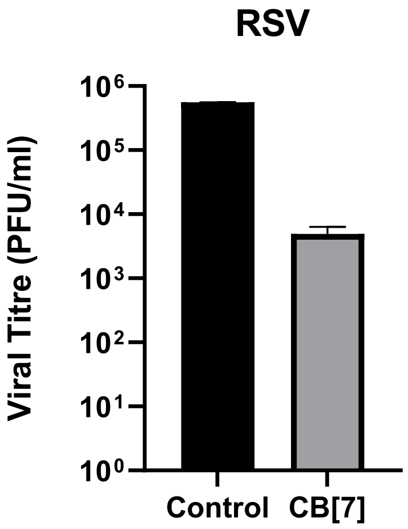


Figure S3: Virucidal assay performed against respiratory syncytial virus (RSV). A virucidal assay is used to distinguish between destructive (virucidal) and non-destructive (virustatic) interactions between antivirals and viruses. In this instance a two log-reduction was observed in RSV titre when 7.5mg/ml of CB[7] was utilised (approximately IC90), indicating a virucidal mode of action.

**TCID$_{50}$ Assay illustrating effect of CBs on MNV1**

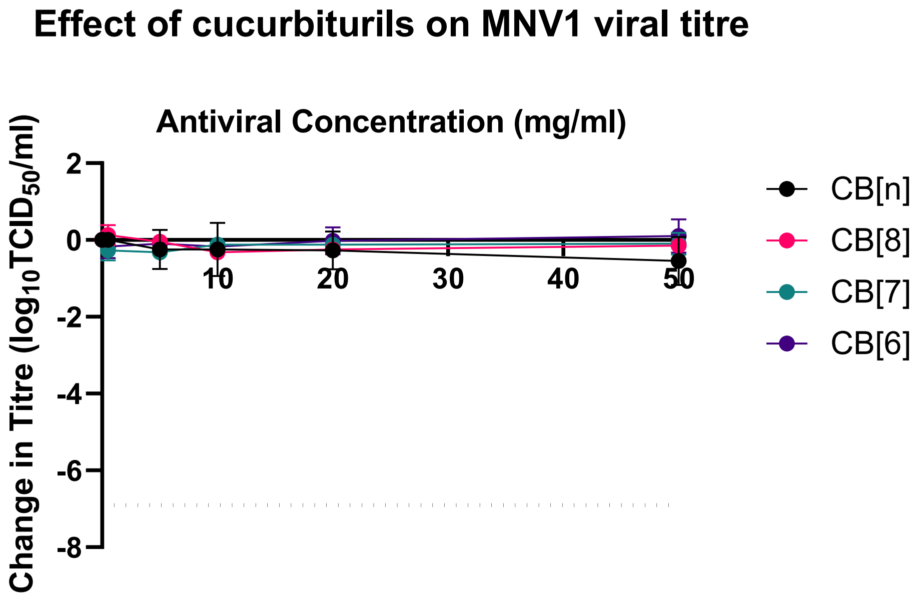


Figure S4: No cucurbituril homologue appeared effective against MNV1 in TCID$_{50}$ assays (n=3 for all homologues).

**Additional Confocal Immunofluorescence Microscopy images for each tested condition of CB[7] with HSV-2**

NTC – Scale Bars = 25 µm

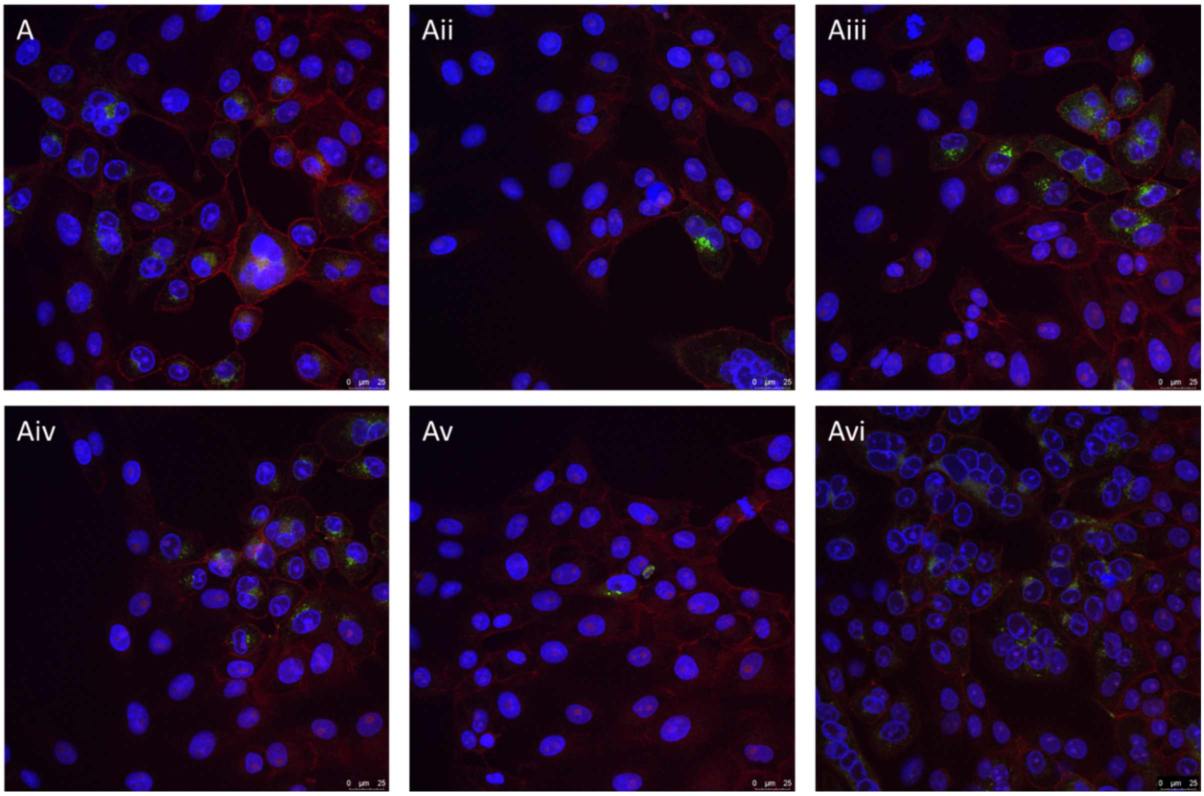


0.05%wt (0.5mg/mlL) CB[7] – Scale Bars = 25µm

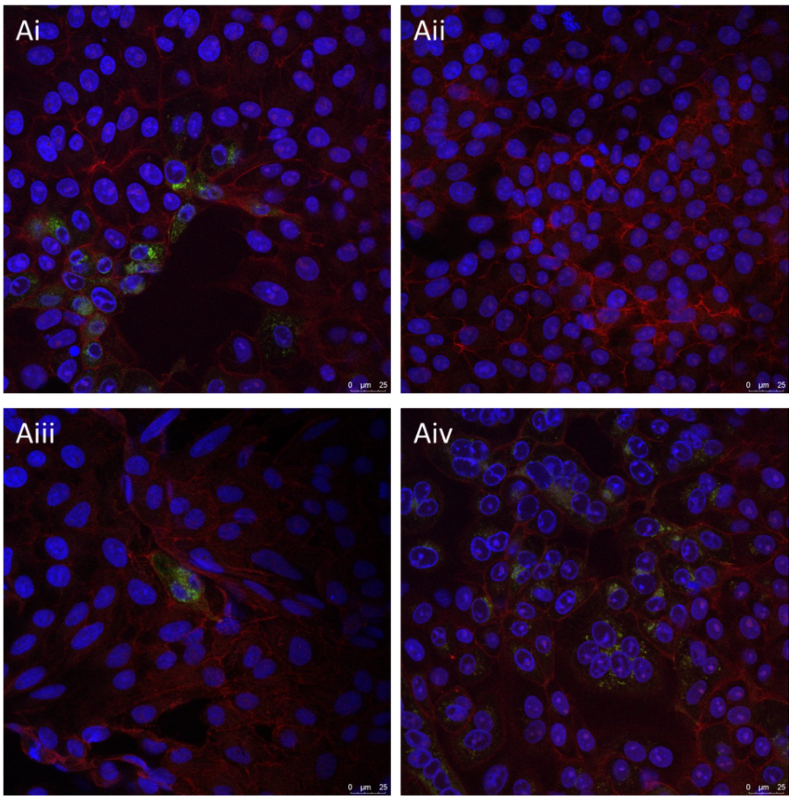

0.5wt% (5mg/mL) CB[*n*] – Scale Bars = 25 µm

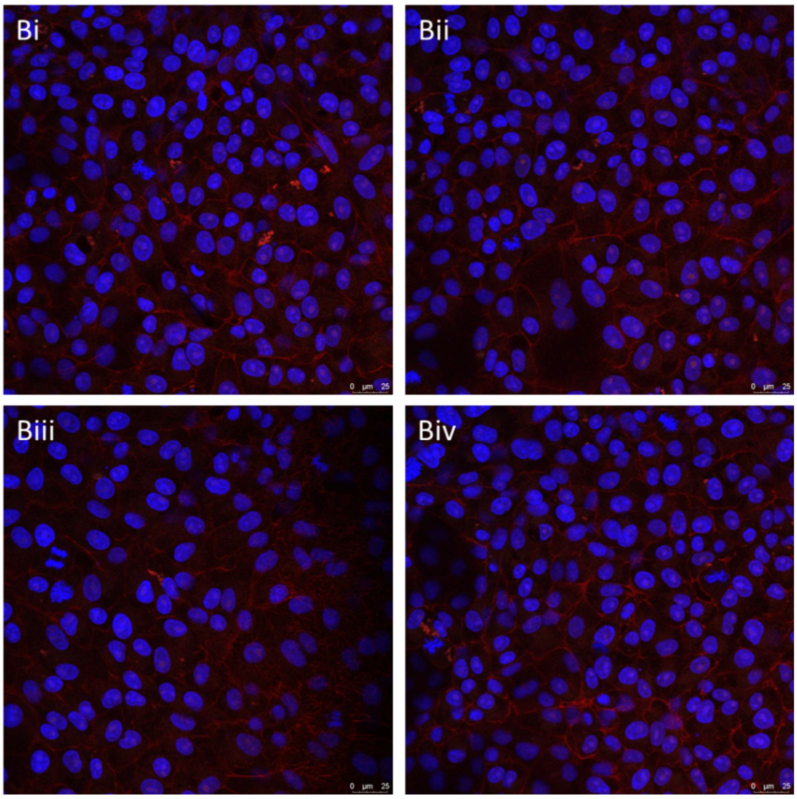


1wt% (10mg/mL) CB[*n*] – Scale Bars = 25 µm

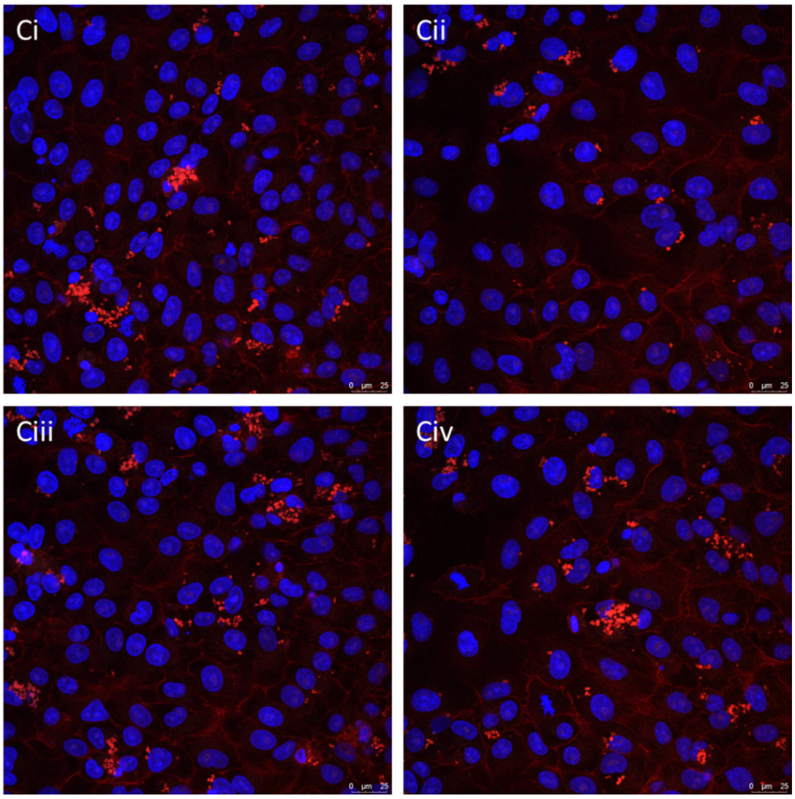

2wt% (20mg/mL) CB[*n*] – Scale Bars = 25 µm

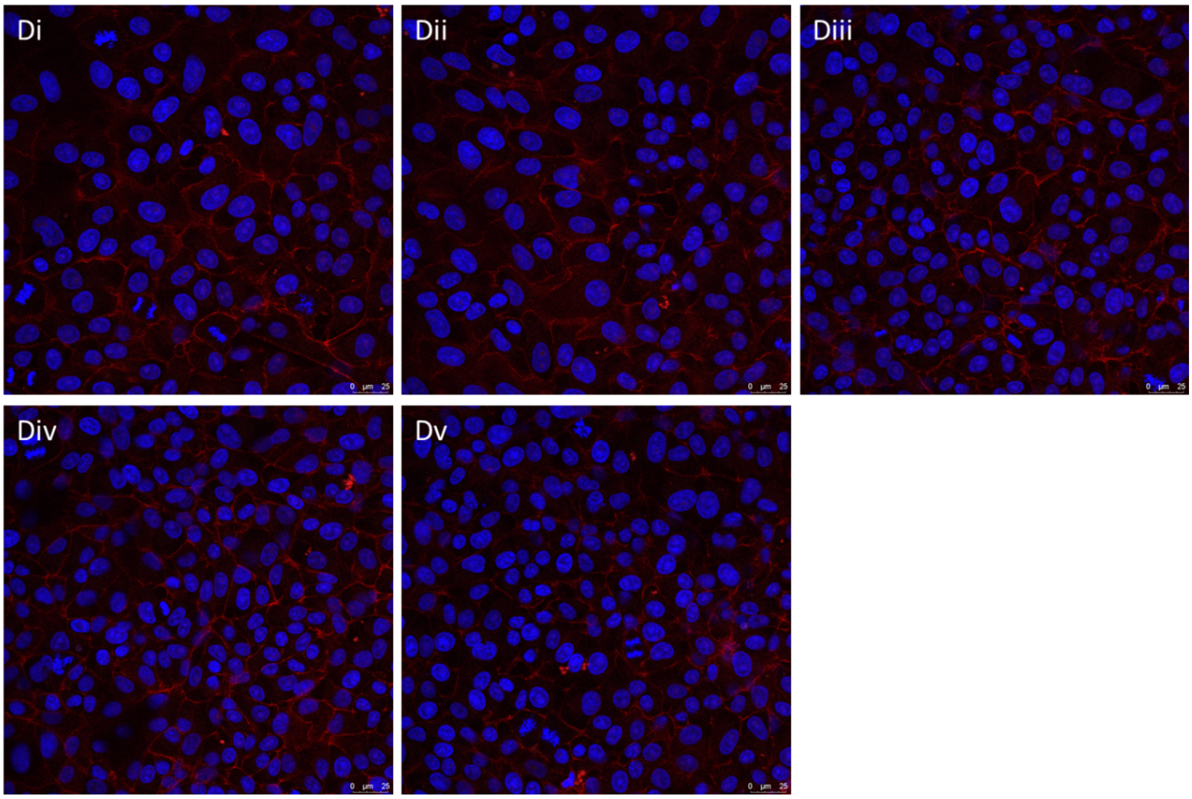


5wt% (50mg/mL) CB[*n*] – Scale Bars = 25 µm

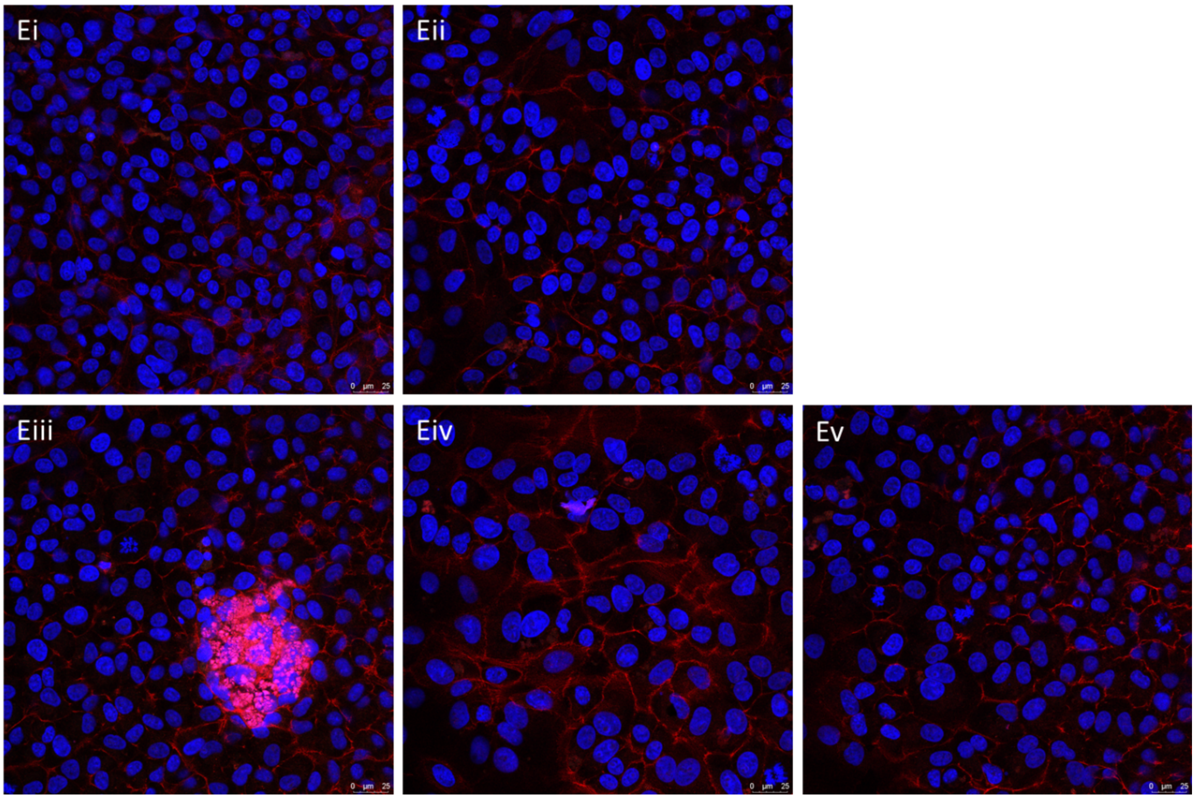

**Additional Confocal Immunofluorescence Microscopy images for each tested condition of CB[*n*] with HSV-2**

NTC – Scale Bars = 25 µm

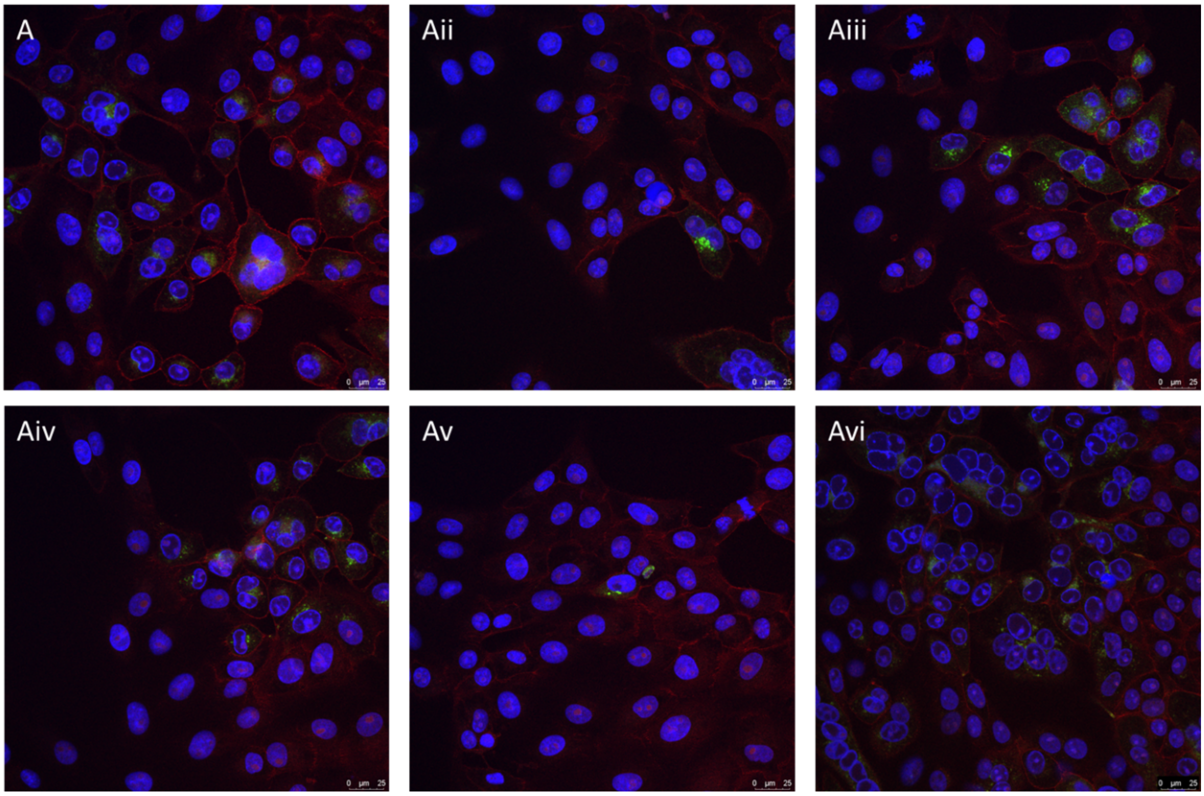


0.5wt% (5mg/mL) CB[*n*] – Scale Bars = 25 µm

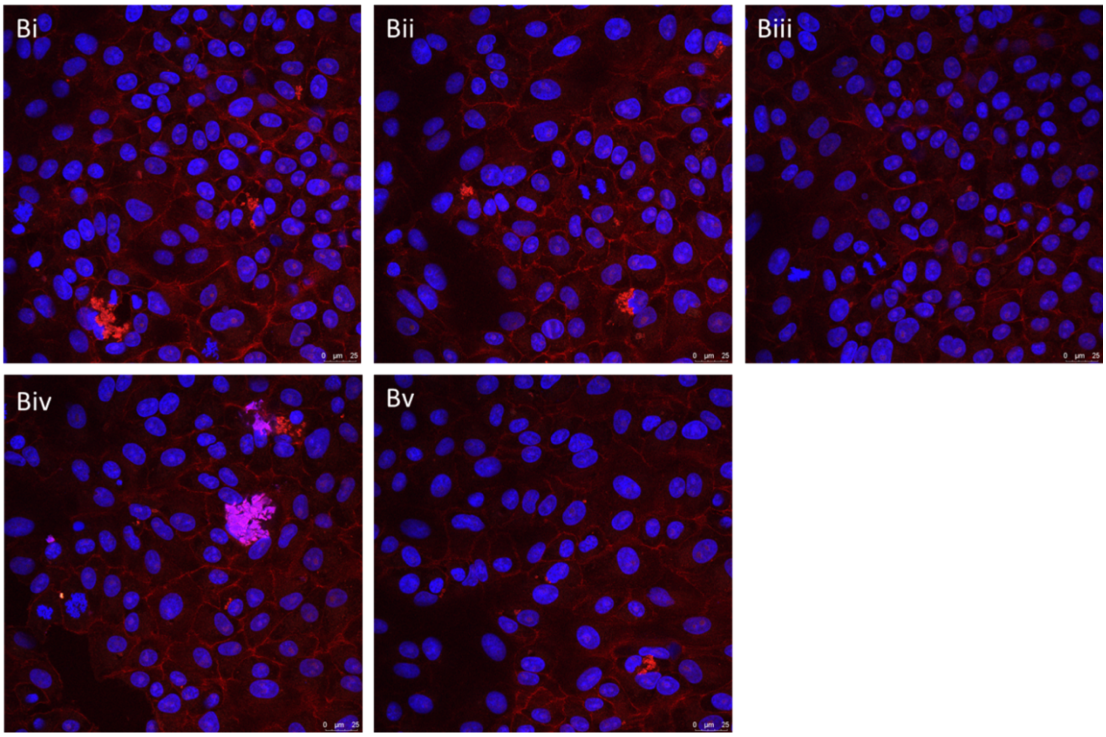

1wt% (10mg/mL) CB[*n*] – Scale Bars = 25 µm

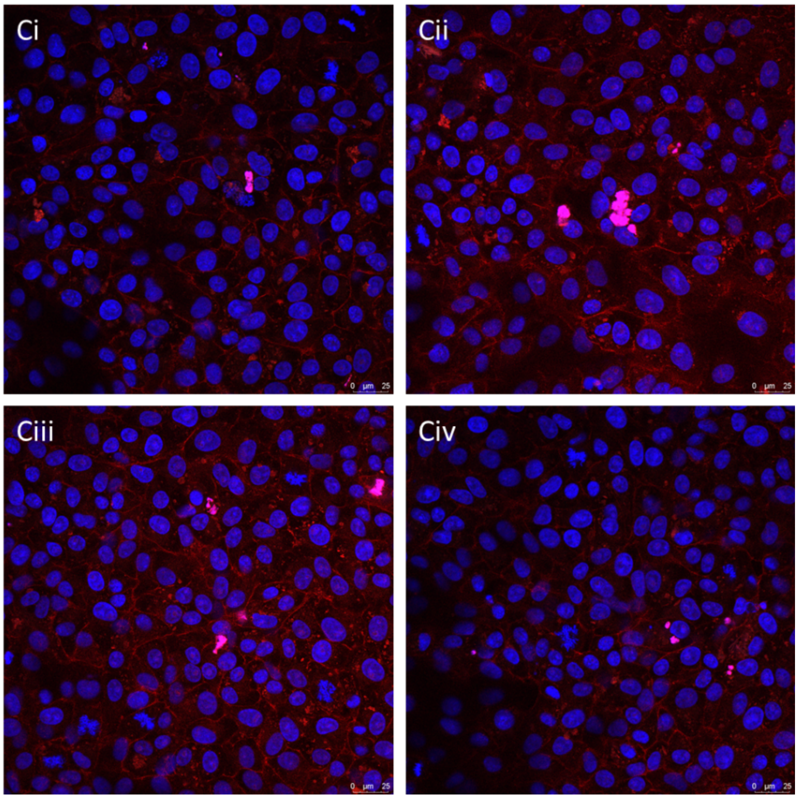


2wt% (20mg/mL) CB[*n*] – Scale Bars = 25 µm

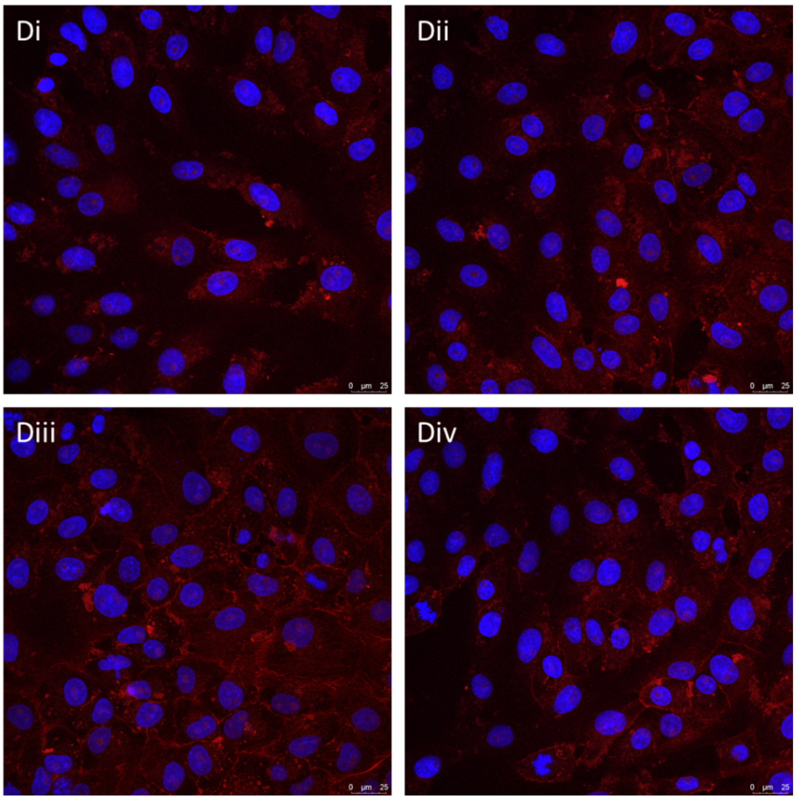

<u>5wt% (50mg/mL) CB[*n*] – Scale Bars = 25 μm</u>

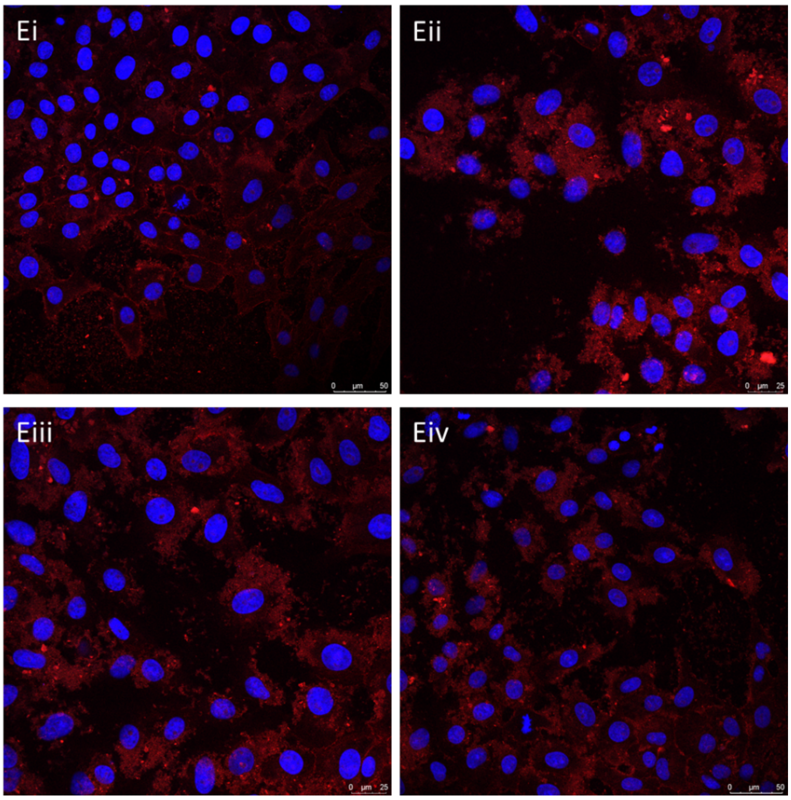
